# Fibrin Polymer on the Surface of Biomaterial Implants Drives the Foreign Body Reaction

**DOI:** 10.1101/2021.02.16.431282

**Authors:** Arnat Balabiyev, Nataly P. Podolnikova, Jacquelyn A. Kilbourne, D. Page Baluch, David Lowry, Azadeh Zare, Robert Ros, Matthew J. Flick, Tatiana P. Ugarova

## Abstract

Implantation of biomaterials and medical devices in the body triggers the foreign body reaction (FBR) which is characterized by macrophage fusion at the implant surface leading to the formation of foreign body giant cells and the development of the fibrous capsule enveloping the implant. While adhesion of macrophages to the surface is an essential step in macrophage fusion and implanted biomaterials are known to rapidly acquire a layer of host proteins, a biological substrate that is responsible for this process *in vivo* is unknown. Here we show that mice with genetically-imposed fibrinogen deficiency display a dramatic reduction of macrophage fusion on implanted biomaterials and are protected from the formation of fibrin-containing granulation tissue, a precursor of the fibrous capsule. Furthermore, macrophage fusion on biomaterials implanted in Fib^AEK^ mice that express a mutated form of fibrinogen incapable of thrombin-mediated polymerization was strongly reduced. Surprisingly, despite the lack of fibrin, the capsule was formed in Fib^AEK^ mice, although it had a different composition and distinct mechanical properties than that in wild-type mice. Specifically, while mononuclear α-SMA-expressing macrophages embedded in the capsule of both strains of mice secreted collagen, the amount of collagen and its density in the tissue of Fib^AEK^ mice was reduced. These data identify fibrin polymer as a key biological substrate driving the development of the FBR.

## INTRODUCTION

Implantation of biomaterials and medical devices in the body triggers the foreign body reaction (FBR) which represents an end-stage of the inflammatory and wound healing responses following injury (1–3). The FBR is characterized by macrophage fusion at the implant surface leading to the formation of multinucleated giant cells, also known as foreign body giant cells (FBGCs) and the development of the fibrous capsule enveloping the implant. FBGC-mediated damage of the biomaterial surface and the formation of a dense fibrous capsule that isolates the implant from the host is the common underlying cause of implant failure.

The very early events following tissue injury caused by implantation include the interaction of blood with the biomaterial surface resulting in adsorption of plasma proteins and the formation of a provisional matrix (1). This matrix serves as an adhesive substrate for neutrophils and monocytes that are recruited out of the vasculature to the implant site with latter cells undergoing differentiation into macrophages. Subsequently, as the acute inflammatory response progresses to chronic inflammation, macrophages at the biomaterial interface fuse to form FBGCs. Macrophage fusion requires both the presence of large foreign surfaces and a microenvironment generated in proximity to the implanted biomaterial. In particular, cytokines IL-4 and IL-13 have been shown to program macrophages into a fusion-competent state *in vitro* (4–6), and IL-4 and IL-13 were identified at the implant site (7, 8, 9). In addition, the chemokine MCP-1 which participates in macrophage fusion *in vitro* and *in vivo* (10) is secreted by biomaterial-adherent macrophages (11).

The formation of macrophage-derived giant cells is concomitant with the growth and remodeling of granulation tissue around the implant which is gradually populated by fibroblasts that transform into myofibroblasts (3, 12). Fibroblasts are thought to be attracted by potent soluble mediators released from activated biomaterial-adherent mononuclear macrophages and FBGCs. Both mononuclear macrophages and FBGCs also secrete pro-fibrogenic factors that enhance fibrogenesis by myofibroblasts (13–15). According to current dogma, the secretion of collagen and other matrix proteins by myofibroblasts results in the production of a fibrous capsule that envelops the implanted device. The capsule containing a dense, avascular layer of collagen is considered an adverse factor of the bioimplant performance because it is impermeable to most molecules in the surrounding environment, thus isolating the implant from the local tissue environment and preventing full healing and incorporation of the implant (16–20). Furthermore, FBGCs themselves are viewed as a significant negative factor contributing to long-term failures in implanted medical devices. FBGCs may cause biomaterial surface damage by releasing potent degradative cellular products such as reactive oxygen intermediates, enzymes, and acid (21–24). Thus, strategies that limit FBGC formation and development of the fibrous capsule are highly desirable.

It is well known that immediately after implantation, biomaterials spontaneously acquire a layer of host proteins that provide an adhesive substrate for arriving inflammatory cells (25, 26). Among them, fibrin(ogen), a principal adsorbed protein, triggers an early arrival and adhesion of phagocytic cells (27–29). Using the intraperitoneal implantation model, Eaton and colleagues have shown that mice made hypofibrinogenemic by injection of ancrod do no mount an acute inflammatory response to the implanted biomaterial (27). Although it has not been determined whether adsorbed fibrinogen or fibrin was responsible for phagocyte recruitment, both proteins can support integrin α_M_β_2_ (Mac-1)- and α_5_β_1_-mediated adhesion of neutrophils and monocyte/macrophages (28, 30–33). Whether fibrinogen or fibrin are also required for macrophage fusion and formation of the fibrous capsule during the FBR is unclear.

In this study, we have shown that macrophage fusion on biomaterials implanted in fibrinogen-deficient (Fg^-/-^) mice was almost completely abrogated and no granulation tissue, a precursor of the fibrous capsule was found around the biomaterials. We further found that Fib^AEK^ mice that express mutated fibrinogen that is incapable of thrombin-mediated polymerization (34) also exhibit a defect in macrophage fusion. Furthermore, although the fibrous capsule in Fib^AEK^ mice was formed and its thickness was comparable to that in wild-type (WT) mice, both matrices had a different composition and distinct mechanical properties. In particular, fibrin-containing capsules formed in WT mice contained greater amounts of collagen than those formed in Fib^AEK^ mice. The cells producing collagen and other extracellular matrix proteins during the early FBR were mononuclear macrophages embedded in granulation tissue. These data indicate that fibrin polymer deposited on the surface of implants is responsible for macrophage fusion and implicate fibrin matrix in organizing a dense fibrous capsule.

## RESULTS

### Formation of FBGCs on implanted biomaterials is abrogated in Fg^-/-^ mice

To assess how fibrinogen might affect macrophage fusion, we used a well-characterized intraperitoneal implantation model to induce the FBR in Fg^-/-^ mice. In these experiments, sterile polychlorotrifluoroethylene (PCTFE) sections were implanted into the peritoneal cavity of WT and Fg^-/-^ mice and the formation of FBGCs was determined after 3, 7, and 14 days. At the time of retrieval, the implanted biomaterial segments were coated by a fibrinous material that appeared as a white film that formed on both sides of the implant in WT mice. Macrophage fusion on the surface of PCTFE implants was determined after the removal of the fibrinous capsule by labeling cells with Alexa Fluor 546-conjugated phalloidin and DAPI. Macrophage fusion in Fg^-/-^ mice was strongly reduced at all time points compared to WT mice (Fig. 1, A and B). On all days, a ∼5-6-fold difference between fusion indices of FBGCs formed in WT and Fg^-/-^ was found. As shown in Fig. 1B, the fusion index in WT mice was increased from 17 ± 8% to 57.0 ± 5% from day 3 to day 14 whereas macrophage fusion in Fg^-/-^ increased from 3.3 ± 2.5% to 9.4 ± 3.9%. The defect in macrophage fusion in Fg^-/-^ mice was not due to the number of macrophages adherent to the implant as even greater numbers of cells (determined as the total number of nuclei) were found on the implants retrieved from Fg^-/-^ mice at days 3 and 7, and equal numbers of cells were found at day 14 (Fig. 1C). Macrophage adhesion and spreading are known to be required for macrophage fusion (35). The degree of mononuclear macrophage spreading to materials implanted into WT mice for 3 days was ∼2-fold greater than in Fg^-/-^ (187 ± 57 vs. 84 ± 37 µm^2^), although it was not significantly different after 7 and 14 days (Fig. 1D). In addition, the migration of leukocytes in Fg^-/-^ mice in response to implantation was not compromised. A trend toward a higher number of cells in the lavage obtained from the peritoneum of Fg^-/-^ mice was noted, but the difference was not significant (Fig. 1E). To examine whether fibrinogen deficiency affects leukocyte migration in another model of inflammation, the leukocyte flux into the peritoneum was induced by the thioglycolate injection. A similar trend was observed, i.e. the total number of monocytes in the peritoneum of Fg^-/-^ mice on day 3 was slightly higher than that in WT mice (Fig. S1). This result is consistent with an increased monocyte/macrophage recruitment into the peritoneum 3 days after thioglycollate injection reported recently by other investigators (36).

**Figure 1.**
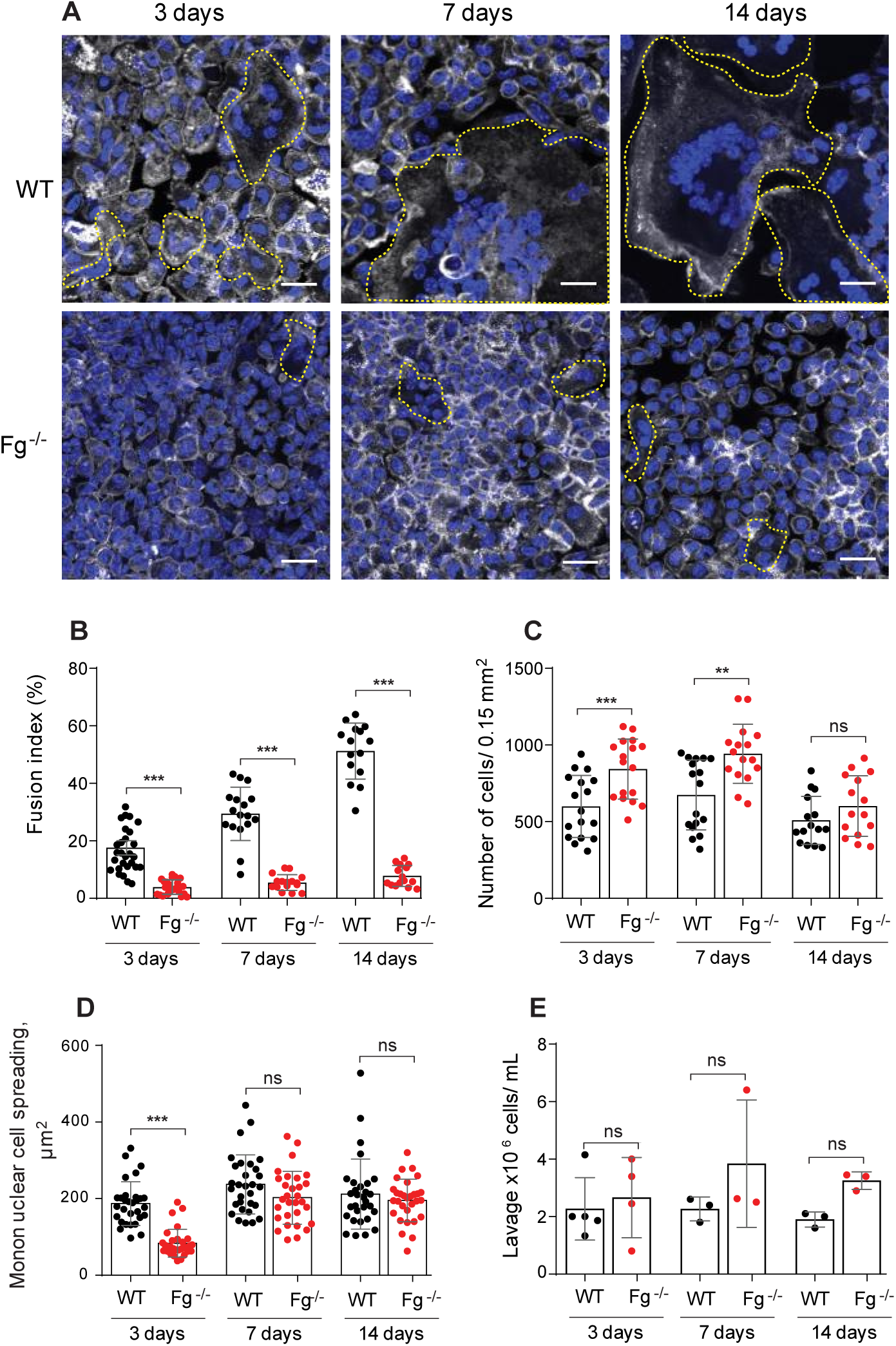
Fg^-/-^ mice display reduced FBGC formation. PCTFE sections were implanted in the peritoneum of wild type and Fg^-/-^ mice for 3, 7, and 14 days and analyzed by immunocytochemistry. (A) Explants were separated from the surrounding fibrous capsule, fixed, and incubated with Alexa Fluor 546-conjugated phalloidin (white) and DAPI (teal). Representative confocal images are shown. FBGCs are outlined (yellow). The scale bar is 20 µm. (B) Macrophage fusion was assessed as a fusion index, which determines the fraction of nuclei within FBGCs expressed as the percentage of the total nuclei counted. Five to six random 20× fields were used per sample to count nuclei. (C) The number of cells on the surface of explants retrieved at various time points was determined by counting nuclei in mononuclear cells and FBGCs. (D) Spreading of mononuclear macrophages on the surface of explants. Six to eight random 20x fields per sample were used to determine the cell area (5-6 cells/field). (E) The number of cells in lavage recovered from the mouse peritoneum before explantation. Results shown are mean ± S.D. of five independent experiments. n=6 (WT) and n=7 (Fg^-/-^) mice per each time point. For counting lavage cells 3-5 WT and 3-4 Fg^-/-^ mice were used per each time point. Two-tailed *t-*test and Mann-Whitney *U* test were used to calculate significance. ns, not significant, \*\**p* < .01, \*\*\**p* < .001.

Histological analyses showed that the thickness of the fibrinous capsule formed around PCTFE material implanted in WT mice gradually increased (Fig. 2, A and B). The capsule was composed of a layer of adherent FBGCs and numerous mononuclear cells embedded into granulation tissue. The capsule was absent around the implants retrieved from Fg^-/-^ mice (Fig. 2, A and B). Even 14 days after implantation, the surface of the implant in Fg^-/-^ mice was only coated with adherent macrophages. Similar results were obtained with polytetrafluoroethylene (PTFE), a biomaterial commonly used for the manufacturing of vascular grafts (Fig. S2). Since PTFE, as compared to PCTFE, is a non-transparent material that precludes direct visualization of macrophages, we used PCTFE in subsequent experiments. Together, these results indicate that macrophage fusion on the surface of the implant and the formation of the granulation tissue, a precursor of the collagenous fibrous capsule, require fibrin(ogen).

**Figure 2.**
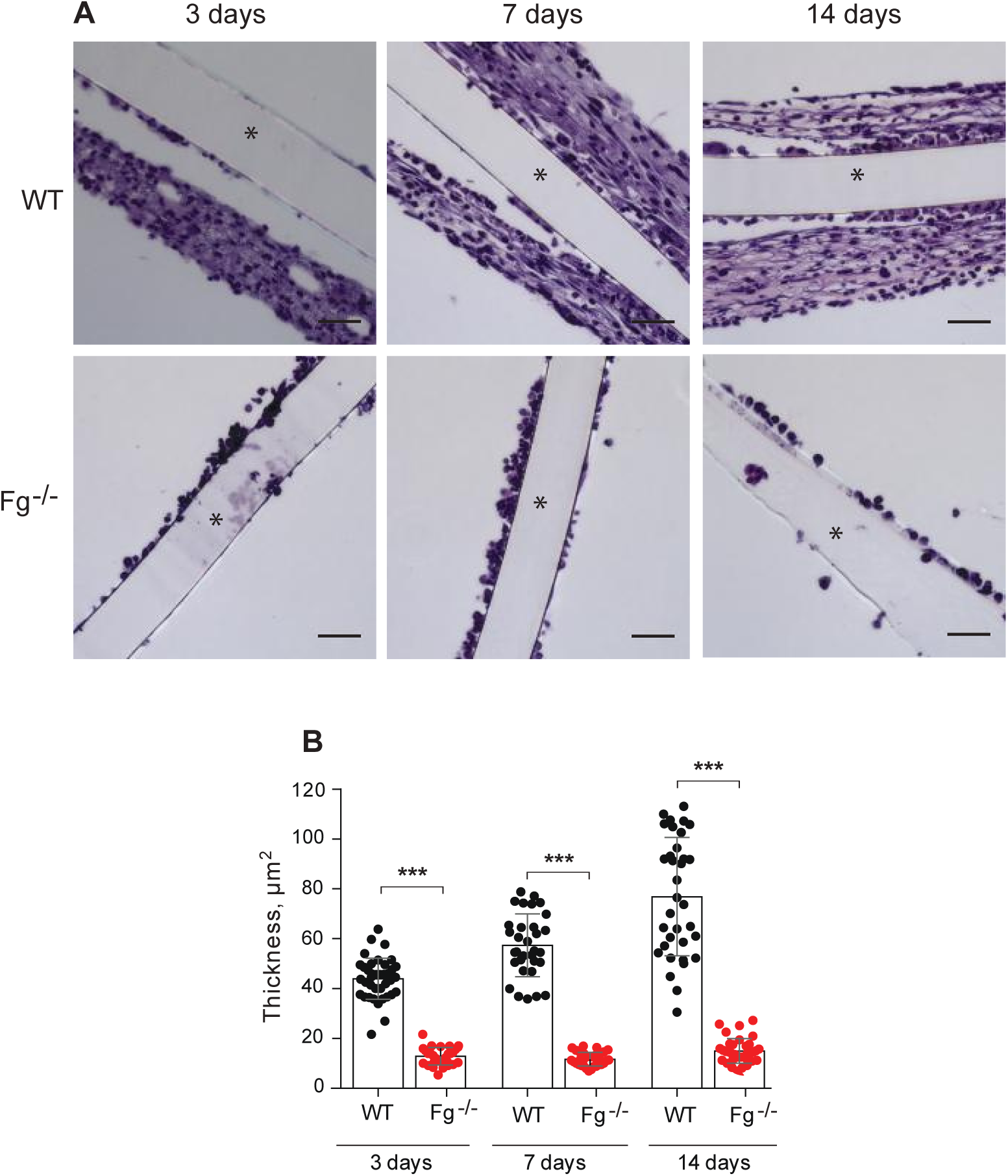
The lack of granulation tissue around the implants in Fg^-/-^ mice. (A) PCTFE sections were implanted in the peritoneum of wild type and Fg^-/-^ mice for 3, 7, and 14 days and analyzed by histochemistry. Explants were fixed, paraffin-embedded, sectioned, and stained according to a standard H&E method. Representative images of stained cross-sections are shown. The scale bar is 50 µm. (B) The thickness of granulation tissue around the implants retrieved from WT and Fg^-/-^ mice was determined using ImageJ software. Ten random fields were used per sample to measure the thickness of cross-sections. Results shown are mean ± S.D. from four independent experiments. n=5 (WT) and n=5 (Fg^-/-^) mice per each time point. \*\*\**p* < .001

### Analyses of fibrinogen and fibrin deposited on the surface of implants and in granulation tissue surrounding the implants

It is generally believed that recruitment and adhesion of leukocytes to the surface of biomaterials implanted into the peritoneum during acute inflammation depends on adsorbed fibrinogen (27). It is possible though that adsorbed fibrinogen is converted into fibrin by thrombin generated at sites of implantation. Therefore, it is not clear whether fibrinogen or the fibrin polymer deposited on the surface of implants mediate macrophage fusion. Furthermore, the fibrino(gen)-rich capsule formed above adherent macrophages in WT mice, which was conspicuously absent in Fg^-/-^ mice, may contribute to macrophage fusion. To investigate these questions, we first examined the presence of fibrinogen and fibrin on the surface of implants and in the capsule (the explant is schematically shown in Fig. 3A; shown is only one side of the explant) using Western blotting with antibodies that recognize the fibrinopeptide A in intact fibrinogen (anti-FpA) and fibrin (anti-Fibrin; mAb 59D8) (Fig. 3B). As expected, the total fibrin(ogen) antigen with a molecular weight of ∼340 kDa corresponding to intact molecule was detected on the surface of implants (Fig. 3C, *left panel*). The lack of reactivity with anti-FpA antibodies indicated that fibrinogen was converted into fibrin (Fig. 3C, *the second panel from the left*). However, no reactivity with fibrin-specific mAb 59D8, which recognizes the N-terminal end of the β-chain in fibrin after cleavage of the fibrinopeptide B (FpB) by thrombin was observed (Fig. 3C, *the third panel from the left*). This finding suggests that fibrin molecules on the surface of implants either lost the epitope for mAb 59D8 due to proteolysis or the epitope is not exposed in the surface-deposited fibrin. A protein band with a molecular weight of ∼150-180 kDa was detected in blots incubated with anti-fibrin mAb 59D8 (*indicated by an arrowhead*). In the absence of the primary antibody, this product interacted with a secondary goat anti-mouse antibody, suggesting that this protein is IgG (Fig. 3C, *right panel*). In line with this finding, previous studies showed that IgG is readily adsorbed from serum on the surface of various materials (37).

**Figure 3.**
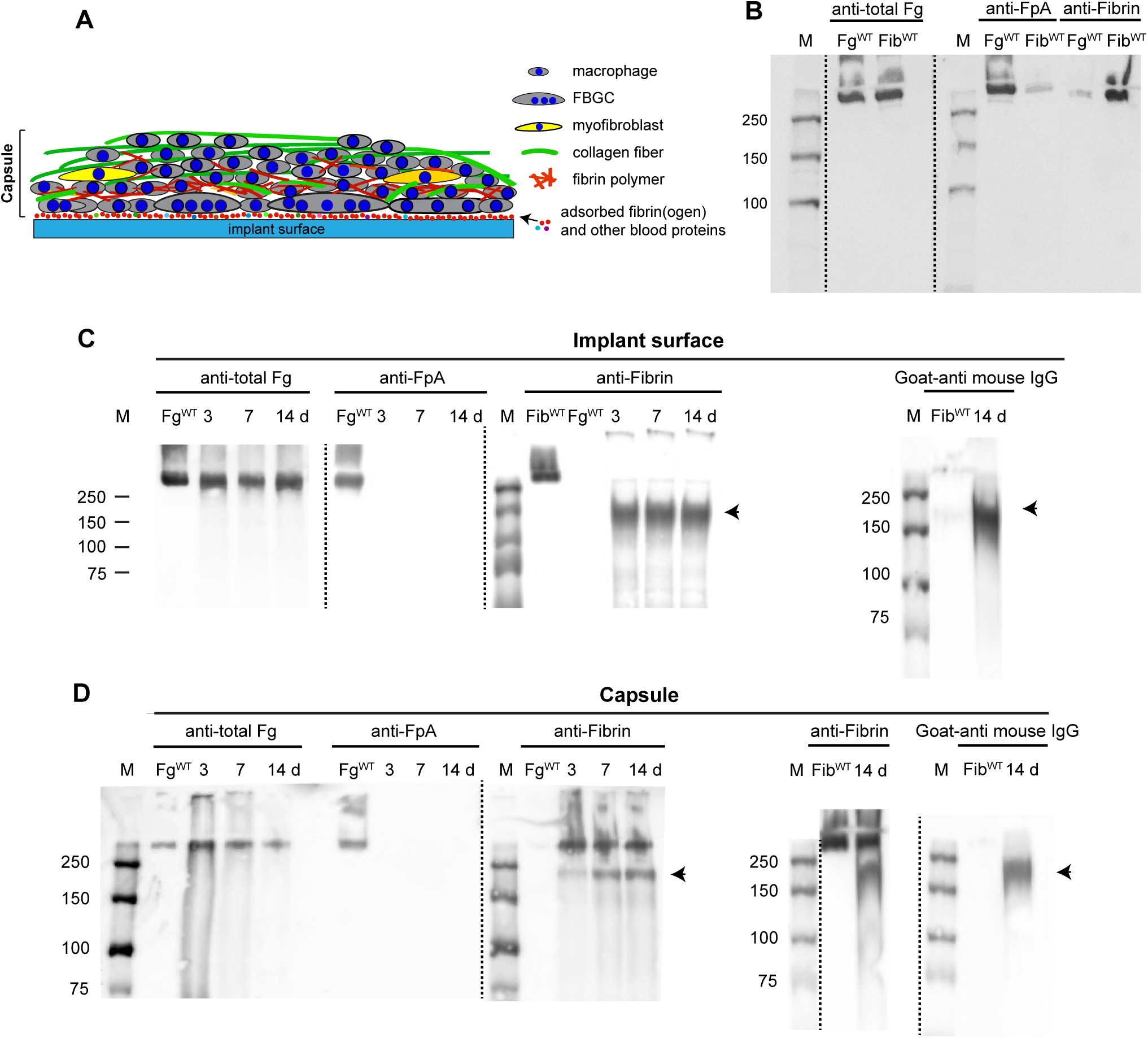
Western blot analysis of proteins deposited on the surface and in the fibrous capsule of materials implanted in WT mice. (A) Schematic representation of proteins and cells present in the explant. The components of the explant are shown on the right. The implants were retrieved after various days and the fibrous capsule was removed from the explant surface. The exposed surfaces with adsorbed proteins and the capsule were analyzed separately. (B) The specificity of antibodies recognizing total fibrinogen (anti-total Fg; 1:50,000 dilution), fibrinopeptide A in intact fibrinogen (anti-FpA; 1:2000 dilution), and the N-terminus of the β-chain in fibrin after the cleavage of fibrinopeptide B (anti-fibrin mAb 59D8; 1 µg/ml). Purified mouse fibrinogen and fibrin-monomer were electrophoresed on 7.5% polyacrylamide gel followed by Western blotting using anti-total fibrinogen, anti-FpA- and fibrin-specific antibodies. (C) Analysis of the implant surfaces retrieved 3, 7, and 14 days after surgery. The sections were placed into PBS containing protease inhibitors followed by the addition of SDS-PAGE loading buffer. The samples were analyzed by Western blotting using anti-total Fg, anti-FpA, and anti-fibrin antibodies. The *right panel* shows the analysis of the material deposited on the surface of the 14-day implant probed with the secondary goat anti-mouse IgG only. The arrowhead indicates a band corresponding to a ∼150-180 kDa product reactive with a secondary goat-anti mouse IgG. The data shown are representative of samples obtained from four mice. (D) Analysis of proteins present in the fibrinous capsule formed at different time points. The right panel shows the analysis of the material obtained from the capsule of the 14-day implant probed with anti-fibrin followed by the secondary antibody or secondary goat anti-mouse IgG only. The IgG-reactive product is indicated by the arrowhead. M, molecular weight markers.

Analyses of the material deposited in the capsule (Fig. 3D) revealed fibrin as evidenced by the presence of the protein with the molecular weight of ∼340 kDa which interacted with mAb 59D8, but not anti-FpA. The IgG-immunoreactive 150-180 kDa protein was also detected in the capsule (Fig. 3D, *right panel*).

### Coating the implant with fibrinogen derivatives rescues the fusion defect in Fg^-/-^ mice

To substantiate the role of fibrin(ogen) in macrophage fusion and determine whether fibrinogen or fibrin was required for this process, materials were coated with purified mouse fibrinogen or fibrin-monomer and implanted in Fg^-/-^ mice for 7 days. Also, to determine whether fibrin-monomer vs. fibrin polymer was responsible for macrophage fusion, a layer of polymerized fibrin gel was deposited on the surface of the material by applying a solution of fibrinogen and thrombin. As shown in Fig.4, A and C, implantation of either fibrinogen- or fibrin-monomer-coated materials in Fg^-/-^ mice restored macrophage fusion by ∼65% of the level observed in WT mice. However, the extent of multinucleation determined as the number of nuclei accumulated in FBGCs, although tended to be higher, was not significantly different from that observed on uncoated surfaces (Fig. 4D). Implantation of sections coated with the fibrin polymer fully restored macrophage fusion (Fig. 4, A and C) and, interestingly, the extent of multinucleation was higher than that in WT mice (Fig. 4D). In addition, pre-coating PCFTE sections with plasma obtained from WT mice and implanted them in Fg^-/-^ mice for 3 days restored macrophage fusion to the level comparable to that in WT mice (15.2 ± 4.6 vs. 18.5 ± 5.7) (Fig. S3, A and C), although the extent of multinucleation was ∼2-fold lower (Fig. S3D). Since all of the explanted surfaces contained a similar number of cells as determined by the number of nuclei, the difference in the rescue effect appears to be due to the form of the fibrin(ogen) substrate and/or its physical properties. Of note, despite the lack of host fibrinogen in Fg^-/-^ mice, a small capsule was formed around materials coated with fibrinogen, fibrin-monomer, and fibrin polymer (Fig. 4B and E) with the capsule around sections coated with fibrin-monomer and fibrin polymer being larger. A small capsule was also formed around the implant coated with plasma (Fig. S3, B and E).

**Figure 4.**
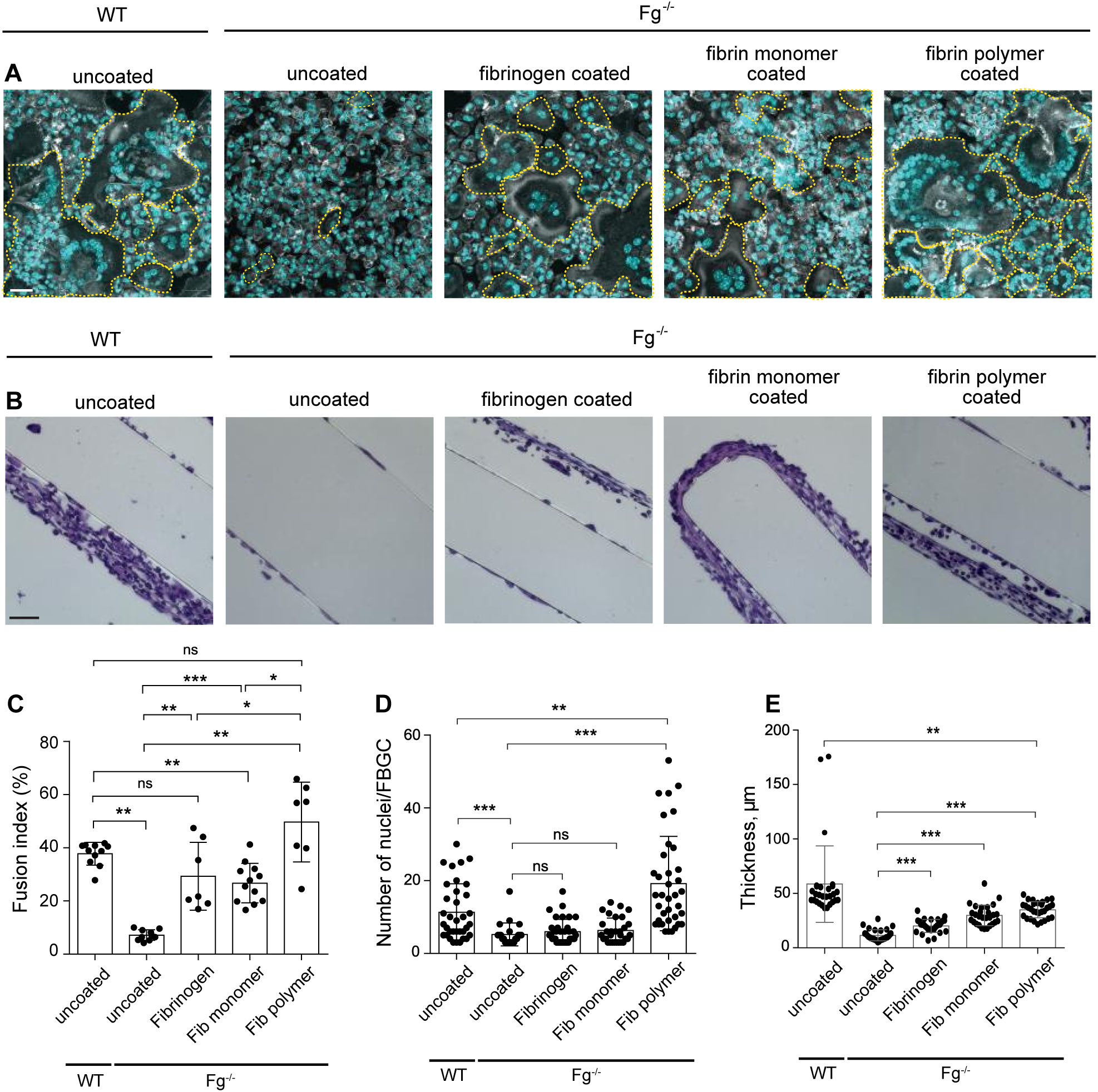
Precoating the implants with fibrinogen and various fibrin(ogen) species restores macrophage fusion on materials implanted in Fg^-/-^ mice. PCFTE sections were precoated with mouse fibrinogen (1 mg/ml), fibrin-monomer (1 mg/ml), or fibrin polymer and implanted into Fg^-/-^ mice for 7 days after which time the implants were removed and analyzed for the presence of FBGCs (A) and granulation tissue formation (B). Uncoated material implanted in WT mice served as a control. (C) Fusion indices were determined as described in Materials and Methods. (D) The extent of multinucleation was determined by counting the number of nuclei per FBGC. ∼70-140 FBGCs were analyzed from 10-15 random fields. (E) The thickness of granulation tissue formed around the plasma-precoated implants was determined using ImageJ software. Ten random fields were used per sample to measure the thickness of cross-sections. Results shown are mean ± S.D. from four independent experiments. n=4 (WT) and n=4 (Fg^-/-^) mice per group. Two-tailed t-test and Mann-Whitney *U* test were used to calculate significance. ns, not significant, *p< .05, **p < .01, ***p < .001. The scale bar is 30 µm in A and 50 µm in B.

To test whether inhibition of fibrin polymer formation could reduce the FBR, we examined the effect of the thrombin inhibitor argatroban on macrophage fusion and the capsule formation in WT mice. Two concentrations of argatroban (9 and 18 mg/kg/day) were injected for 5 days before implantation of biomaterials and then daily for 7 days post-surgery, after which materials were explanted and analyzed. In a pattern analogous to that observed for implants in Fg^-/-^ mice, treatment of WT mice with argatroban significantly reduced macrophage fusion compared to untreated controls (∼1.8- and 2-fold for 9 and 18 mg/kg, respectively; Fig, 5A and 5C), even though a slightly larger or similar number of cells was present on the surface of materials implanted in treated mice (Fig. S4). FBGCs on the surface of implants in argatroban-treated mice contained fewer nuclei (Fig. 5, D and E) and were smaller (Fig. 5F and G). However, the thickness of the capsule formed around materials in treated and untreated mice was not significantly different (Fig. 5, B and H). Together, these data suggest that fibrin-polymer deposited on the surface of implants is required for macrophage fusion but may be dispensable for capsule formation.

**Figure 5.**
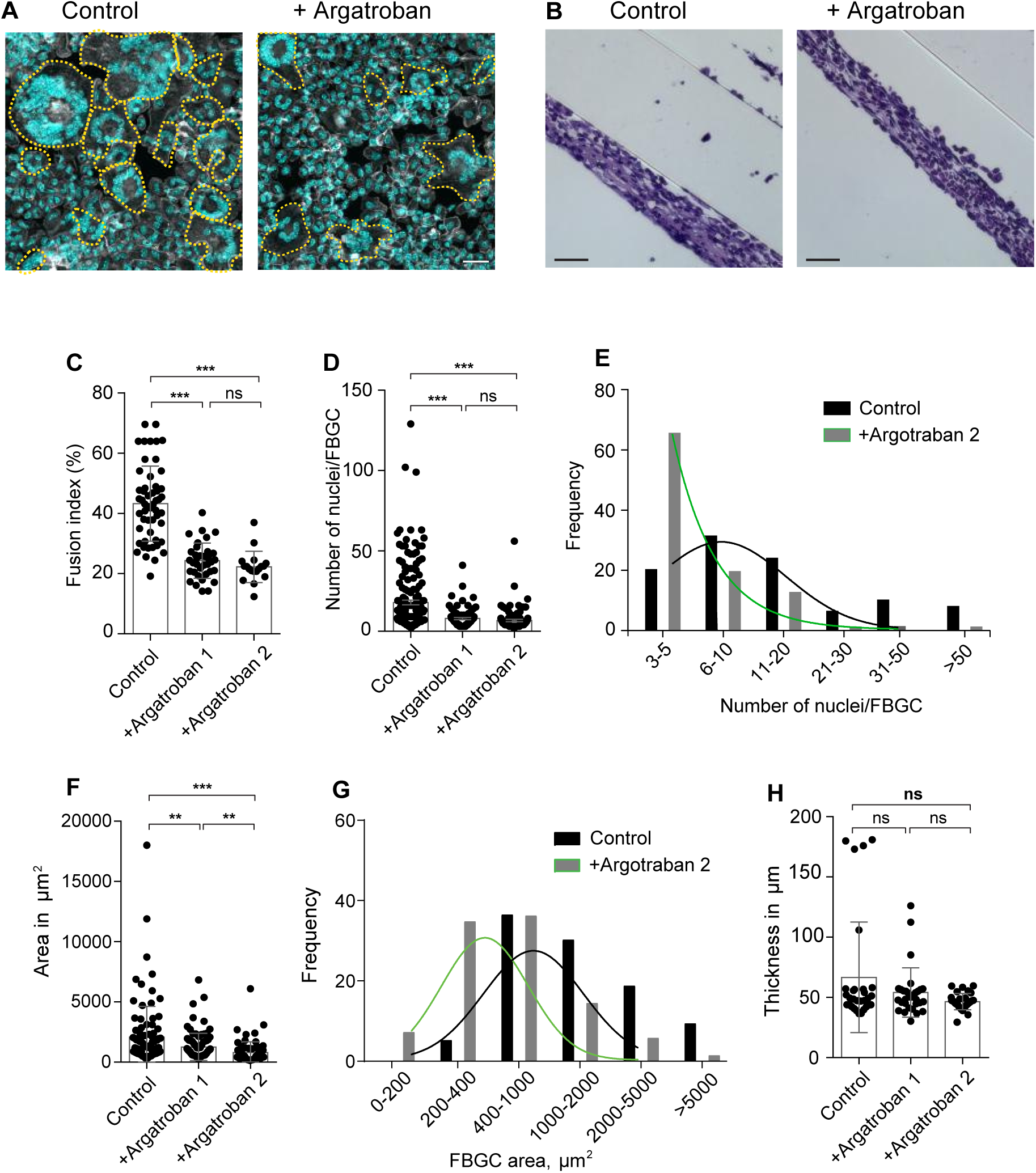
Thrombin inhibitor reduces macrophage fusion but does not affect capsule formation. (A) Representative confocal images showing that treatment with argatroban, as compared with the untreated control, reduces FBGC formation. Argatroban (9 mg/kg and 18 mg/kg; termed argatroban 1 and 2, respectively) was injected i.p. in WT mice for 5 days before implantation of materials and continued for 7 days post-surgery. Control mice were injected with PBS. (B) Granulation tissue formed around implants was stained with H&E. (C) Macrophage fusion was assessed as a fusion index. Five to six random 20× fields were used per sample to count nuclei. (D) The number of nuclei in FBGCs on the surface of implants retrieved from control and argatroban-treated mice. Approximately 70-140 FBGCs on each implant was counted. (E) The frequency distribution of nuclei in FBGCs on implants retrieved from control mice and mice treated with 18 mg/kg argatroban. (F) Spreading of FBGCs on the surface of explants retrieved from control mice and mice treated with 18mg/kg argatroban. Six to eight random 20x fields per sample were used to determine the cell surface area (10-15 cells/field). (G) The frequency distribution of the FBGC areas on implants retrieved from control mice and argatroban-treated mice. (H) The thickness of granulation tissue formed on the surface of implants retrieved from control and argatroban-treated mice was determined using ImageJ software. Ten random fields were used per sample to measure the thickness of cross-sections. Results shown are mean ± S.D. from three independent experiments. n=6 mice per group. Two-tailed *t-*test and Mann-Whitney *U* test were used to calculate significance. ns, not significant, \*\**p* < .01, \*\*\**p* < .001

### Macrophage fusion on materials implanted in Fib^AEK^ mice carrying a mutation in the thrombin-cleavage site in the A**α** chain of fibrinogen is compromised

To directly determine the role of fibrin polymer in mediating macrophage fusion, PCTFE sections were implanted in Fib^AEK^ mice and various parameters of the FBR were determined. In these mice, the six Aα chain amino acid residues upstream of the thrombin cleavage site (Glu^P6^-Gly-Gly-Gly-Val-Arg^P1^) were mutated to prevent the removal of FpA thus incapacitating the ability of fibrinogen to polymerize (34). Similar to the FBR observed in Fg^-/-^ mice, macrophage fusion was impaired (***Fig. 6A and 6C***) with the fusion index being ∼6-fold lower at day 14 in Fib^AEK^ mice compared to WT mice. Furthermore, similar to Fg^-/-^ mice, the reduced fusion in Fib^AEK^ mice was not due to a decrease in the number of adherent macrophages as comparable (day 3) or even greater numbers of cells (days 7 and 14) were found on the surface of implants (***Fig. 6D***). However, in contrast to Fg^-/-^ mice, where a difference between mononuclear macrophage spreading on implanted surfaces was observed only at day 3, cell spreading on surfaces retrieved from Fib^AEK^ mice was significantly reduced at all times (***Fig. 6E***). No significant difference between the number of leukocytes in the lavage of WT and Fib^AEK^ mice was found (***Fig. S5***).

**Figure 6.**
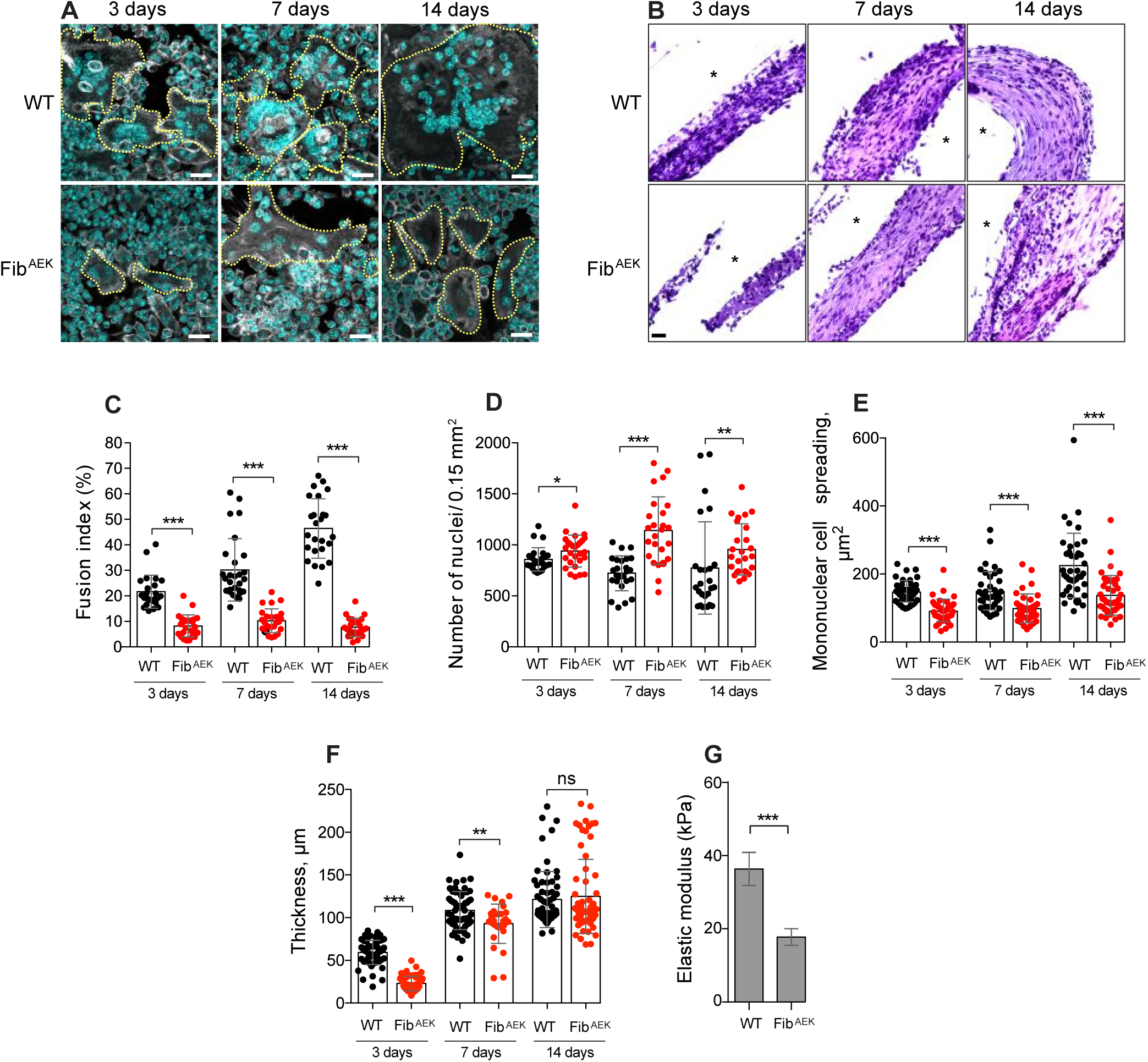
The formation of FBGCs and not the production of the capsule is compromised in Fib^AEK^ mice. PCTFE sections were implanted in WT and Fib^AEK^ mice for 3, 7, and 14 days. (A) Explants retrieved after selected time points were separated from the surrounding fibrinous capsule, fixed, and incubated with Alexa Fluor 546-conjugated phalloidin (white) and DAPI (teal). Representative confocal images are shown. The scale bar is 20 µm. (B) Representative images of the H&E stained cross-sections of implants retrieved from the peritoneum of WT and Fib^AEK^ mice The scale bar is 20 µm. (C) Fusion indices of macrophages formed on the surface of implants that have been retrieved at different time points from WT and Fib^AEK^ mice. (D) The density of cells on the surface of explants retrieved at various time points was determined by counting the number of nuclei. (E) Spreading of the mononuclear cells on the surface of implants. (F) The thickness of the capsule formed in WT and Fib^AEK^ mice. Results shown are mean ± S.D. of four independent experiments. n=4 (WT) and n=4 (Fib^AEK^) mice. Two-tailed *t-*test and Mann-Whitney *U* test were used to calculate significance. ns, not significant, \**p <* .05, \*\**p* < .01, \*\*\**p* < .001. (G) Mechanical properties of the 14-day capsules formed around implants retrieved from WT and Fib^AEK^ mice were determined by measuring the elastic moduli (expressed in Pa) using AFM. Results are mean and S.E. of 300 force-indentation curves on each sample

Western blot analyses of fibrinogen species deposited on the surface of implants demonstrated the availability of total fibrinogen antigen with a molecular weight of 340 kDa (Fig. S6B, *left panel*). No reactivity with anti-FpA was detected (Fig. S6B, *middle panel*). The lack of reactivity with anti-FpA mAb can potentially arise from the substitution of six amino acid residues in the FpA rather than the cleavage of the peptide. Indeed, fibrinogen isolated from Fib^AEK^ mice failed to interact with the anti-FpA antibody (Fig. S6A, *right panel*). Moreover, consistent with failed fibrin polymerization, we were unable to produce fibrin from fibrinogen isolated from Fib^AEK^ mice and consequently detect the interaction of anti-fibrin mAb 59D8 with this protein. Hence, no interaction of mAb 59D8 with proteins deposited on the surface of implants was found (Fig. S6B, *right panel*). Similar to WT mice, the presence of a product with a molecular weight of ∼150-180 kDa reactive with a secondary goat anti-mouse antibody was observed.

Surprisingly, although less robust on day 3, granulation tissue-like material was formed around PCTFE sections implanted in Fib^AEK^ mice and its thickness was comparable to that in WT mice at days 7 and 14 (Fig. 6, B and F). Western blotting showed that the material present in the capsule contained fibrinogen as evidenced by the interaction with an antibody directed against the total protein and an IgG-immunoreactive 150-180 kDa product (Fig. S6C).

Grossly, the capsules retrieved from WT and Fib^AEK^ mice were indistinguishable but Fib^AEK^-derived material was softer and more fragile, indicating that the mechanical properties of these matrices were different. To quantitatively assess the difference, we determined the stiffness of each tissue using AFM. The analysis showed that the elastic modulus of granulation tissue isolated from WT mice was 2.3-times greater than that of tissue retrieved from Fib^AEK^ mice (37.0 ± 2.2 kPa vs. 16.2 ± 0.6 kPa; Fig. 6G).

### Analysis of the fibrous capsule formed around implants in WT and Fib^AEK^ mice

Since the capsule was formed around implants in Fib^AEK^ mice in the absence of fibrin polymerization we sought to determine its composition. Because fibrinogen was detected by Western blotting, we first examined its spatial distribution within the capsule and compared it with the capsule retrieved from WT mice using immunohistochemistry. As shown in Fig. 7A and 7B, the extensive deposition of fibrinogen and fibrin were detected in the capsule retrieved from WT mice. Examination of the capsule in Fib^AEK^ mice revealed the presence of total fibrinogen antigen that increased by day 14 (Fig. 7C). Consistent with the results of Western blotting, no fibrin was detected (Fig. 7D).

**Figure 7.**
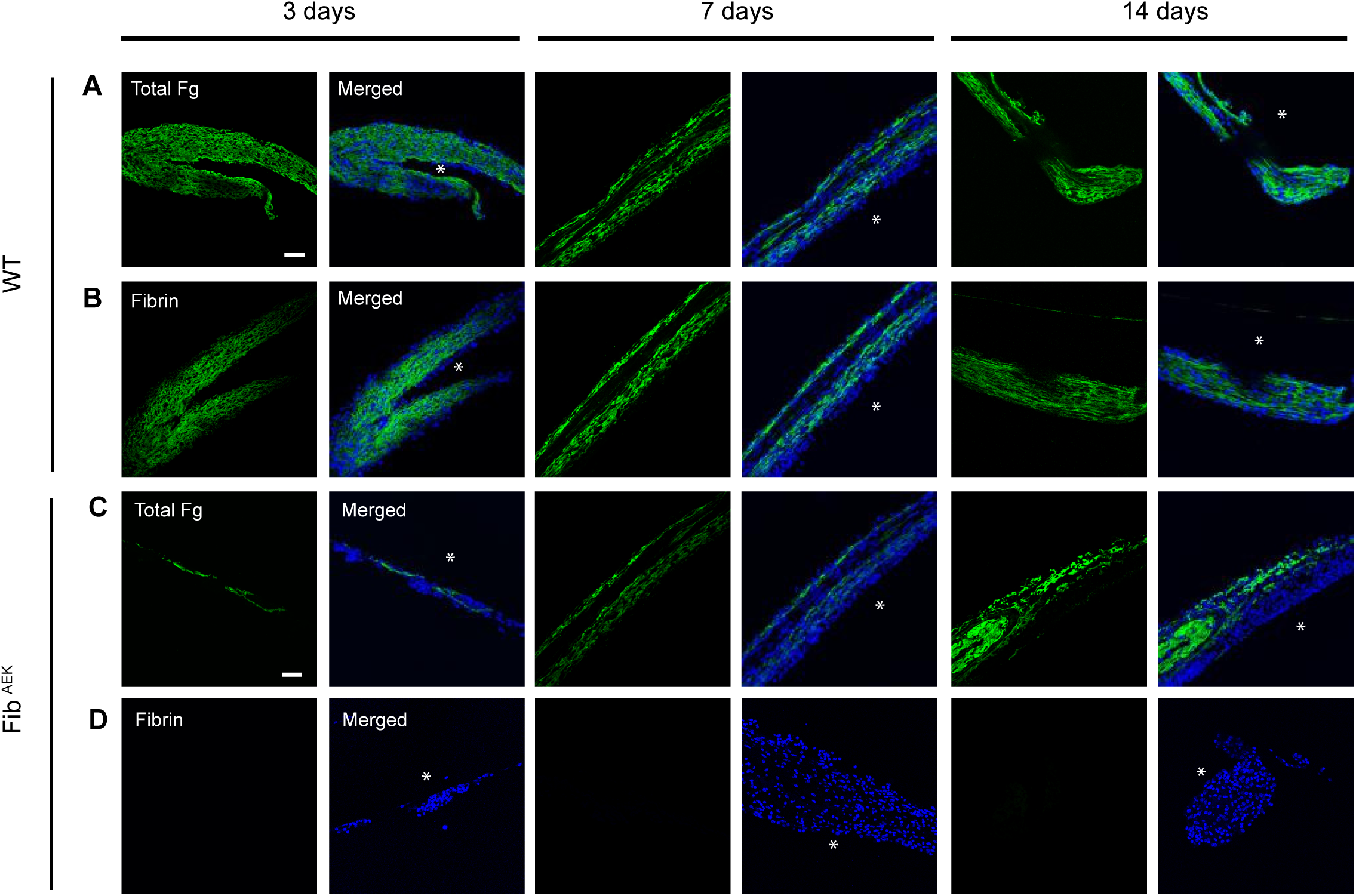
Distribution of fibrinogen and fibrin in the capsule formed on the surface of implants in WT and Fib^AEK^ mice. Representative confocal images of tissue formed on the surface of implants 3, 7, and 14 days after surgery. The samples were incubated with antibodies that recognize total fibrinogen (A, C) and fibrin (B, D). The scale bar is 50 µm.

Next, we determined whether other proteins may organize the capsule in Fib^AEK^ mice. Since collagen is a component of the fibrous capsule, we examined the time-dependent deposition of collagen in both WT and Fib^AEK^ mice using a mAb directed to Type 1α collagen. The collagen density in the capsule was quantified by measuring the green-pixel coverage per 400 µm^2^ area. While no collagen was detected in the 3-day capsule formed in both strains of mice, it was detectable on day 7 and its amount strongly increased on day 14 in WT mice (Fig. 8, A and C). In comparison, the amount of collagen was significantly less in the capsule from Fib^AEK^ mice (Fig. 7, B and C). Additional investigations of the capsule organization using transmission electron microscopy (TEM) confirmed the presence of abundant collagen fibrils in WT mice that were organized into fibers that ran approximately at right angles to one another (Fig. 8D, shown for day 14). Collagen fibrils were also observed in the capsule of Fib^AEK^ mice (Fig. 8D); however, fibrils were significantly shorter than in WT mice (590±343 nm vs. 1515±697 nm) (Fig. 8E). Moreover, the density of fibrils determined as the distance between individual fibrils was significantly greater in WT- than in Fib^AEK^-derived tissue (18.2±3.9 nm vs. 27.2±4.5 nm) (Fig. 8F). We also found that material retrieved from the WT and Fib^AEK^ capsules contained fibrillin and elastin, typical components of the extracellular matrix of connective tissue (Fig. S8, A-D). These proteins were first detected as early as day 3 and their amounts increased by day 14. The density of fibrillin and elastin in the capsule from WT mice was greater than in Fib^AEK^ mice on day 3 and was not significantly different at later time points (Fig. S8, E and F).

**Figure 8.**
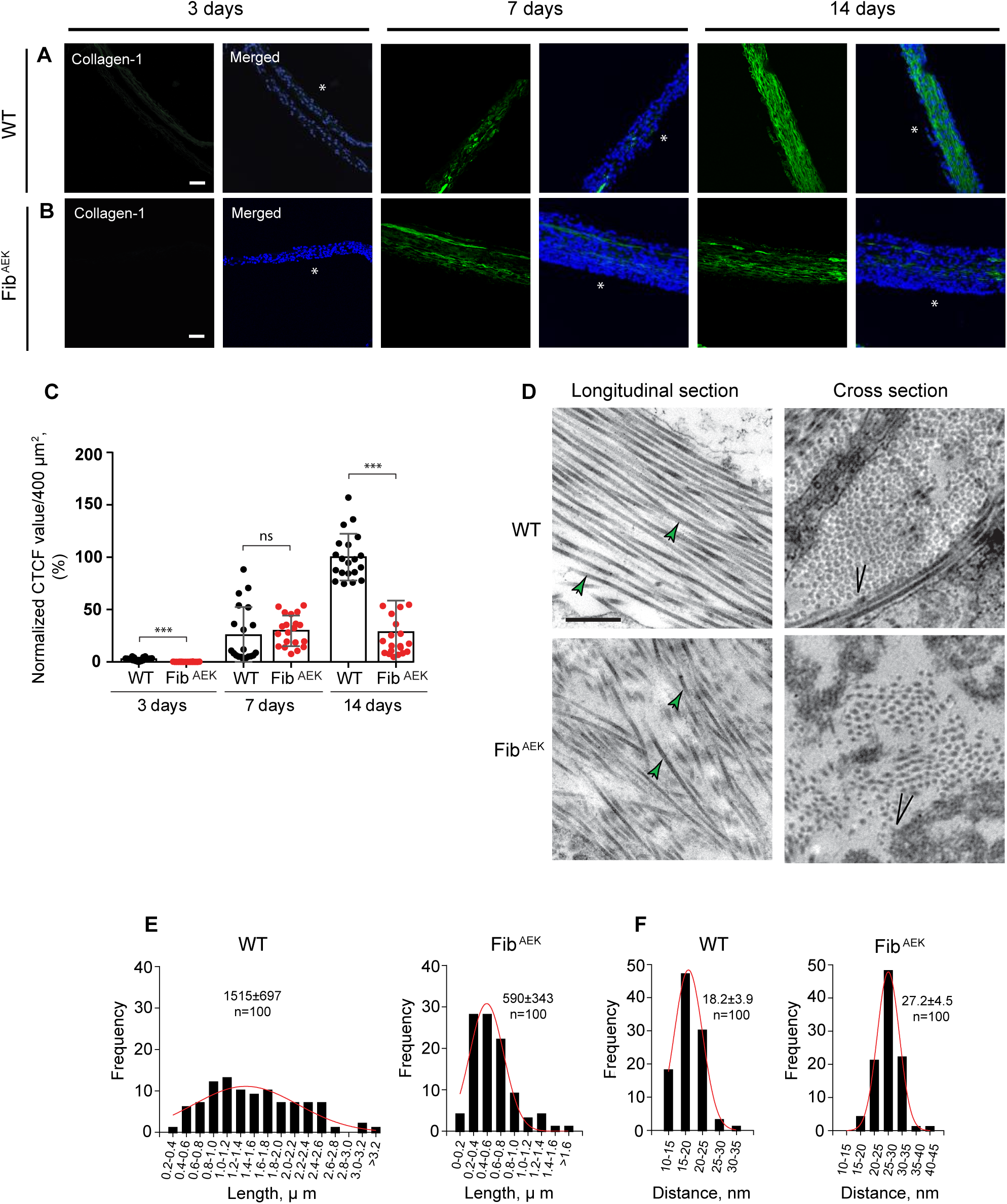
Deposition of collagen in the capsule formed on the surface of implants in WT and Fib^AEK^ mice. PCTFE sections were implanted in WT and Fib^AEK^ mice for 3, 7, and 14 days and the capsules formed were analyzed for the presence of collagen. (A, B) Representative immunofluorescence images of the samples incubated with antibodies that recognize collagen I. The scale bars are 50 µm. (C) Quantification of fluorescence intensities of images of collagen deposition shown in A and B. CTCF (correlated total cell fluorescence) values were determined by ImageJ software using the following formula: Integrated Density - (Area of selected region x Mean fluorescence of background readings). Collagen expression in the 14-day capsule was assigned a value of 100%. Results shown are mean ± SD of three independent experiments. n=3 (WT) and n=3 (Fib^AEK^) mice. Mann-Whitney *U* test was used to calculate significance. ns, not significant, \*\*\**p* < .001. (D) The ultrastructural details of the capsule were examined by TEM. Representative TEM micrographs of the sections prepared by cutting the specimen parallel (*longitudinal)* and vertical (*cross)* to the substratum are shown. The scale bar is 500 nm. (E) The frequency distribution of the collagen fiber lengths in the capsule from WT and Fib^AEK^ mice. (F) The frequency distribution of the collagen fiber densities in the capsule from WT and Fib^AEK^ mice.

### Mononuclear macrophages in the capsule express collagen

It is generally believed that mononuclear macrophages and FBGCs recruit fibroblasts that begin to invade granulation tissue 2 - 3 weeks after implantation, differentiate into myofibroblasts, and initiate collagen deposition (16, 38). The absence of cells with morphological features of fibroblasts/myofibroblasts in the TEM images and rather an early deposition of collagen in the capsule from WT and Fib^AEK^ mice suggested that other cells may secrete collagen. To identify these cells, we labeled the material in the capsule retrieved from WT and Fib^AEK^ mice with a mAb recognizing α-smooth muscle actin (α-SMA), a myofibroblast marker, and CD68, a macrophage marker. Many cells in the capsules retrieved at 3-14 days appeared to show the presence of both proteins (Fig. 9, A-D). However, the tight packing of cells in the capsule precluded the quantification of cells that expressed only α-SMA, only CD68, or both proteins. Therefore, we isolated cells from the 14-day capsules and analyzed them by immunocytochemistry using mAbs directed to α-SMA and CD68. The analyses showed that a small population of cells (∼6%) expressed α-SMA and were negative for CD68 (Fig. 10, A and B; shown for cells isolated from a 14-day WT capsule). Based on these features and the presence of actin stress fibers, these cells were identified as myofibroblasts (Fig. 10 A, *upper panel*). The main population of cells (∼ 93%) isolated from the WT and Fib^AEK^ capsules expressed α-SMA and CD68, displayed podosomes, and were thus identified as monocyte/macrophages (Fig. 10, A and B). MAb M1/70 which recognizes myeloid-specific CD11b/CD18 (integrin Mac-1) also labeled these cells (Fig. 10C, *bottom panel*) but not myofibroblasts (Fig. 10C; *upper panel*). Both myofibroblasts and monocyte/macrophages were stained for collagen using mAb directed against collagen 1α (Fig. 10C). Collagen expression was confirmed using polyclonal anti-collagen I antibody (Fig. S8A). Interestingly, collagen expression in WT macrophages was ∼2.4-fold greater than in their Fib^AEK^ counterparts (Fig. 10D). To exclude the possibility that staining for collagen was due to the uptake by macrophages of collagen secreted by fibroblasts, we performed RT-PCR experiments using purified macrophages. The analyses demonstrated the presence of mRNA for collagen I (Fig. S8B). Expression of fibrillin and elastin in macrophages isolated from the capsule was also detected; however, expression of these proteins did not differ between WT and Fib^AEK^ mice (Fig. S9). Together, these results suggest that cells of monocyte origin present in the capsule are the main source of collagen production and that WT monocytes secrete more collagen than cells accumulated in the Fib^AEK^ capsule.

**Figure 9.**
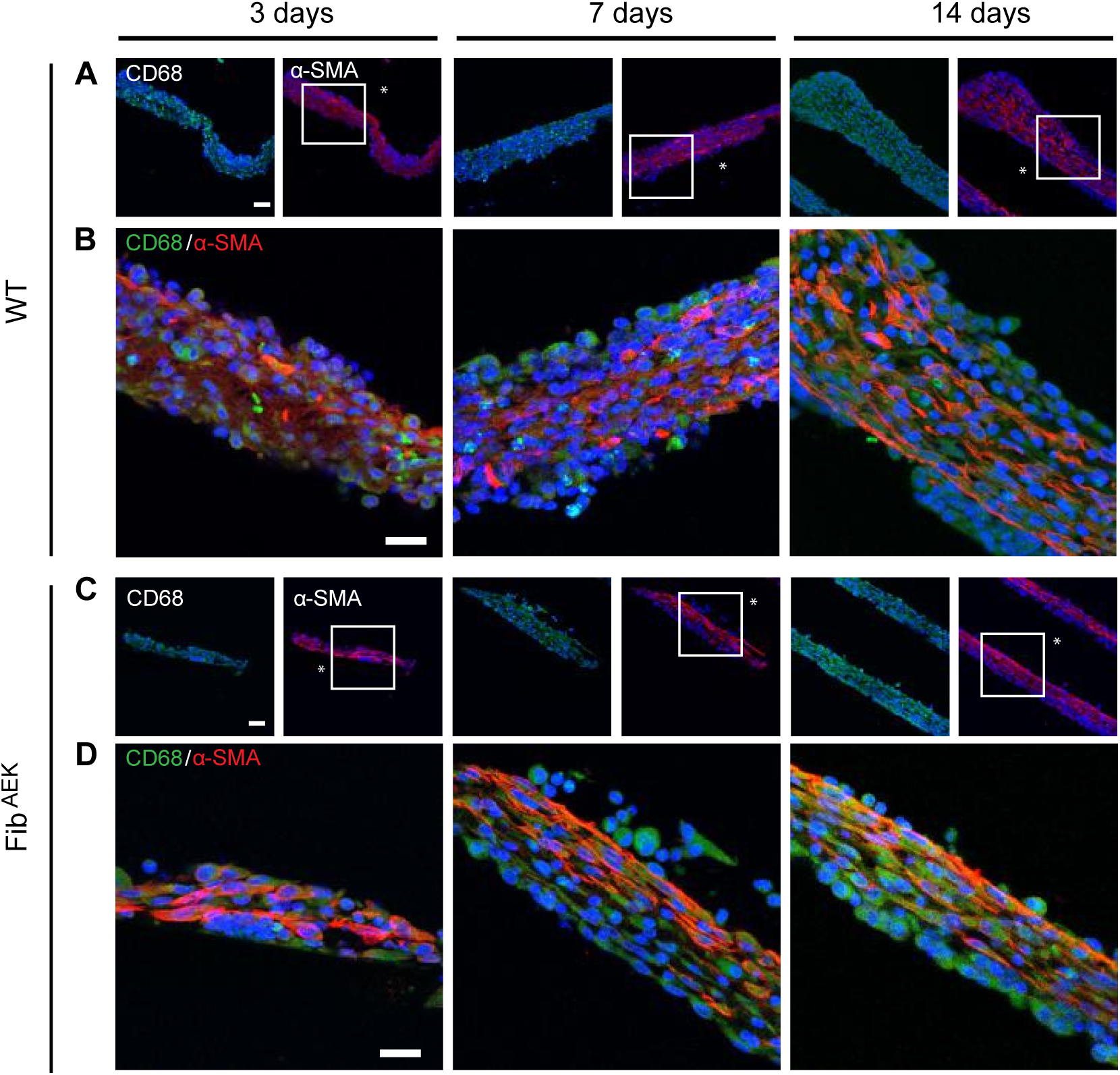
Macrophages within the fibrous capsule express α-SMA and CD68. PCTFE sections were implanted in WT (A, B) and Fib^AEK^ (C, D) mice for 3, 7, and 14 days. Representative images of the capsule samples retrieved from WT (n=3) and Fib^AEK^ (n=3) mice incubated with anti-CD68 (green) and anti-α-SMA (red) antibodies are shown. High magnification views of boxed areas in A and C are shown in B and D. The scale bars are 50 μm (A and C) and 20 μm (B and D).

**Figure 10.**
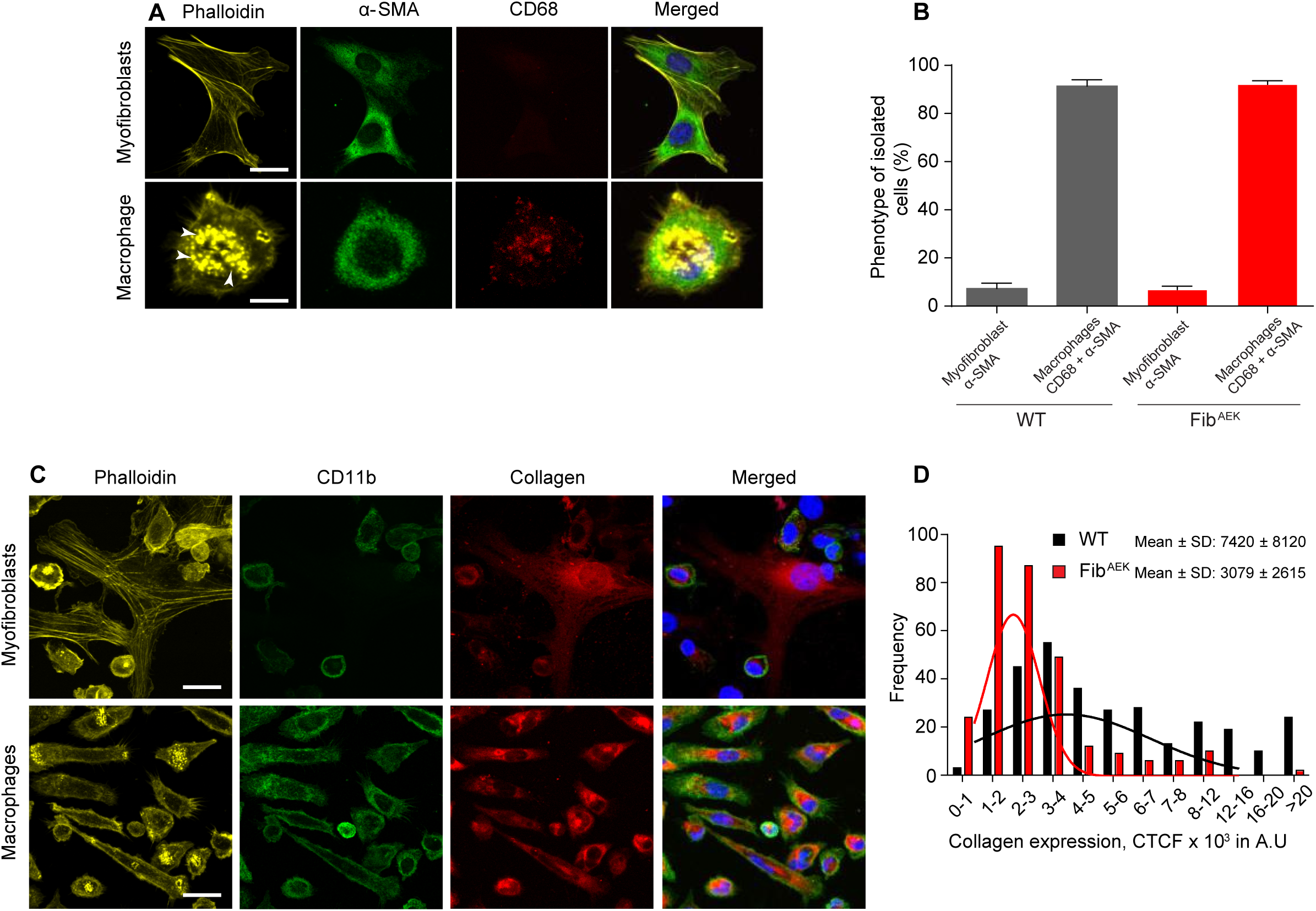
Macrophages within the fibrous capsule express collagen. PCTFE sections were implanted in WT and Fib^AEK^ mice for 14 days and the cells accumulated within the capsules were isolated as described in the Materials and Methods. (A) The cells were allowed to adhere to the surface of a FluoroDish, fixed, and incubated with Alexa Fluor 568-conjugated phalloidin, anti-α-SMA, and anti-CD68 antibodies. Representative confocal images of myofibroblasts and macrophages isolated from the capsule retrieved from WT mice are shown. Arrowheads point to podosomes seen in macrophages incubated with phalloidin-Alexa Fluor 568. The scale bar is 10 μm. (B) Quantification of cells expressing αSMA only (myofibroblasts) or both αSMA and CD68 (macrophages). Approximately 2000-2500 cells isolated from the capsules retrieved from each WT and Fib^AEK^ mice were analyzed with ∼400 cells present in each field of view. (C) The cells isolated from the capsule obtained from WT mice were incubated with anti-CD11b mAb M1/70 and anticollagen I mAb. The scale bar is 20 μm. (D) The frequency distribution of fluorescence intensities for collagen I in macrophages isolated from the capsules retrieved from WT and Fib^AEK^ mice was expressed as CTCF arbitrary units (A.U). A total of 300 cells from 5 random fields in the samples prepared from WT and Fg^AEK^ mice were analyzed. The results shown are from 3 independent experiments. n=3 (WT) and n=3 (Fib^AEK^) mice.

## DISCUSSION

The foreign body reaction (FBR) to implanted biomaterials begins with the spontaneous adsorption of host proteins within seconds after contact with body fluids (1). This process initiates the recruitment and accumulation of phagocytes on the surface of implants. Among adsorbed proteins, fibrin(ogen) is chiefly responsible for this early inflammatory response (27). In the present study, we have examined the role of fibrino(gen) in mediating the later stages of FBR, including macrophage fusion and fibrous capsule formation. Several lines of evidence indicate that fibrin polymer deposited on the surface of implants is required for macrophage fusion. First, the presence of fibrinogen, a fibrin precursor, is indispensable for the formation of macrophage-derived FBGCs since the implantation of materials into the peritoneum of Fg^-/-^ mice resulted in the almost complete lack of macrophage fusion. However, despite the presence of intact fibrinogen in Fib^AEK^ mice that express a mutated form of fibrinogen incapable of thrombin-mediated polymerization, macrophage fusion was also strongly reduced. Second, implantation in Fg^-/-^ mice of materials coated with fibrin polymer fully rescued the fusion defect whereas coating with fibrinogen or fibrin-monomer partially normalized macrophage fusion. Third, inhibition of thrombin with argatroban reduced macrophage fusion. These findings indicate that fibrin polymer is a major determinant of FBGCs formation.

Previous studies conducted in mice made hypofibrinogenemic by injections of ancrod, a thrombin-like protease from the venom of Malayan pit viper showed almost no phagocyte accumulation on materials implanted for 16 hours in the mouse peritoneum and this defect could be normalized by coating materials with fibrinogen or by injection of purified fibrinogen (27). In addition, studies conducted in Fg^-/-^ mice showed a ∼ two-fold decrease in the number of phagocytes adherent to implanted biomaterial 18 hours after implant (39). Our data showed that 3 days after implantation, when macrophage fusion begins, the surfaces implanted in WT and Fg^-/-^ mice contain similar numbers of mononuclear cells (Fig. 1C), suggesting that at later times, adsorption of other plasma proteins can support phagocyte adhesion. However, only the absence of fibrin(ogen) in Fg^-/-^ mice resulted in the dramatic defect in macrophage fusion. Moreover, analyses of the surfaces explanted from WT mice showed that adsorbed fibrinogen converted into fibrin as was evidenced by the presence of the product with the molecular weight of ∼340 kDa that lacked FpA (Fig. 3B). Curiously, fibrin deposited on the surface did not interact with mAb 59D8 which recognizes the N-terminal portion of the β-chain of fibrinogen after cleavage of FpB. This finding can be explained by the propensity of the β-chain to undergo rapid degradation by various proteases releasing the β15-41 and β42-53 fragments (40, 41) or by the fact that the epitope is not exposed in the surface-deposited fibrin.

Consistent with the requirement for fibrin, coating of materials with fibrin polymer normalized macrophage fusion in Fg^-/-^ mice. Importantly, since the coating of implants with purified fibrinogen or fibrin-monomer only partially restored macrophage fusion, the polymeric state of fibrin seems to be essential for effective fusion. The requirement for fibrin polymer rather than adsorbed fibrinogen in mediating macrophage fusion is puzzling since it is generally believed that adsorbed fibrinogen, due to its partial unfolding, has altered conformation and shares many properties with fibrin, including expression of the binding sites for integrins, monoclonal antibodies, and other molecules (42, 43). The fusion-promoting effect of the fibrin network deposited on the surface of implants may be indirect and involve trapping of specific cytokines and chemokines involved in the induction of the fusion program. Alternatively, the physical properties of the fibrin matrix *versus* adsorbed fibrinogen can potentially influence macrophage fusion by initiating a specific mechanotransduction response. In this regard, numerous studies demonstrated that the physical properties of extracellular matrices, including those made of fibrin(ogen) influence intracellular signaling and cell responses (44–47). Indeed, we have observed that mononuclear cell spreading, a sign of integrin-mediated signaling, and the ensuing actin cytoskeleton rearrangement was lower on the surfaces implanted in Fib^AEK^ mice compared to WT mice (Fig. 6E). The actin cytoskeleton is known to be actively involved in macrophage fusion (48–50) and thus the interaction of macrophage integrins with fibrin matrices may initiate alterations of the actin cytoskeleton conducive to fusion. Yet another possibility is that fibrin polymerization results in the exposure of sequences that are not present in the soluble and adsorbed forms of fibrinogen and which may serve as binding sites for cellular structures involved in macrophage fusion. This proposal is based on a precedent established by studies showing that the binding of t-PA and VE-cadherin is mediated by fibrin-specific sequences, not accessible in fibrinogen (51–53). At this juncture, the basis for the differential effect of fibrinogen and fibrin remains speculative and further studies may help to define the mechanisms underlying the fusion-promoting activity of fibrin polymer.

Our data show that fibrinogen is required for the formation of granulation tissue and, ultimately, the fibrous capsule surrounding the implant. Only a layer of adherent cells but a complete lack of the capsule was detected around materials implanted in Fg^-/-^ mice (Fig. 2). At the same time, although fibrin polymerization was compromised, the formation of the capsule in Fib^AEK^ mice was not impaired, suggesting that fibrinogen was required for the capsule formation (Fig. 6F). Furthermore, a small capsule was formed around the materials coated with fibrinogen, fibrin-monomer, and fibrin polymer that were implanted into Fg^-/-^ mice (Fig. 4). In agreement with these data, biomaterials coated with polyethylene oxide-like compounds that are known to be ultra-low fouling and inhibit fibrinogen adsorption (54) reduced the fibrous capsule thickness (55). Analogously, zwitterionic materials with ultralow-fouling properties (56) resisted the capsule formation for several months after subcutaneous implantation (57). Therefore, the deposition of fibrin(ogen) on the surface of implants seems to nucleate the fibrous capsule formation.

Analyses of the capsule formed in WT and Fib^AEK^ mice showed that in addition to fibrinogen it contains collagen and other proteins typically present in the extracellular matrix of connective tissue, including fibrillin and elastin. However, the amount of collagen in the capsule from WT mice was significantly higher and collagen fibers were longer and packed into denser bundles than in the capsule from Fib^AEK^ mice. These morphological features were reflected in the different mechanical properties of both matrices. Based upon the measurement of the elastic modulus, the matrices formed in WT mice were about 2.3-fold stiffer than those in Fib^AEK^ mice. These findings suggest that fibrin polymer abundantly deposited in the capsule of WT mice may organize collagen and other proteins into a dense matrix. This is, to the best of our knowledge, the first evidence for a critical role of fibrin polymer in the organization of the fibrous capsule.

It is generally believed that cells responsible for collagen production and deposition in the fibrous capsule are α-SMA-expressing myofibroblasts. Our data show that cells producing collagen I and other connective tissue proteins indeed express α-SMA. However, these cells also express CD68 and CD11b/CD18 (integrin Mac-1), suggesting that these cells are of monocyte/macrophage origin. This finding is corroborated by the fact that collagen I was deposited in the capsule between 3-7 days and fibrillin and elastin were detected as early as 3 days after implantation, i.e. much faster than would be expected for the secretion of these proteins by recruited fibroblast/myofibroblasts. Indeed, myofibroblasts were identified only in a 14-day capsule and their proportion was small (∼6%) (Fig. 10B). Given this small number, it seems unlikely that the deposition of collagen in the capsule was due solely to myofibroblasts.

Recent murine and human studies provided evidence that monocyte-derived cells at sites of injury express α-SMA and secrete collagen I (58–60). These cells originate from a subset of circulating monocytes that express collagen I, a mesenchymal marker, and CD45 and CD11b/CD18, hematopoietic markers (61–63). Cells that combine a gene expression profile of macrophages with that found in fibroblasts and have both the inflammatory features of macrophages and the tissue remodeling properties of fibroblasts have been termed fibrocytes (58, 59, 61). Fibrocytes participate in both physiological wound healing and pathological fibrosis (58, 59). Furthermore, fibrocyte-like cells have been detected in the vicinity of biomaterials (64, 65). Mooney and colleagues showed that cells within the capsule formed in response to implantation in the peritoneal cavity of a foreign object express α-SMA protein (65) and also express genes typically found in macrophages (66). It is presently unclear whether α-SMA^+^/CD68^+^/CD11b^+^/Col^+^ cells infiltrating the capsule in our implant model are recruited from blood or originate from other sources. Although further studies are needed to elucidate the phenotype of these cells and their origin, this is the first demonstration that cells expressing both hematopoietic and stromal markers are involved in the deposition of collagen and other extracellular matrix proteins in the capsule during the FBR.

Previous studies revealed many aspects of macrophage fusion, including the requirement for adhesion, the induction of specific intracellular signaling, and the role of specific molecules (35). The majority of this information was generated *in vitro* experiments using macrophages cultured on various surfaces, including biomaterials, with only a few studies performed *in vivo* models of implantation (10, 67–71). Moreover, with the exception of plasma fibronectin (pFN), little consideration has been given to proteins that may serve as adhesive substrates for fusing macrophages *in vivo.* Using pFN conditional knock-out mice, Keselowsky *et al* demonstrated that the number of FBGCs on biomaterials implanted subcutaneously was three times higher than in WT mice (72). The mechanism by which pFN exerts this effect remains elusive and the role of cellular fibronectin which was not depleted needs to be investigated. McNally *et al* showed that among several plasma proteins that might adsorb on the surface of implants, vitronectin mediated the greatest IL-4-induced fusion of cultured monocyte/macrophages (73). While vitronectin can serve as important adhesive ligands mediating initial macrophage adhesion to the surface of implants, its role in macrophage fusion has not been corroborated in the present study as the deficiency of a single protein, fibrinogen, completely abrogated macrophage fusion. More importantly, studies conducted in a unique Fib^AEK^ mouse model allowed us to conclude that it is fibrin rather than intact fibrinogen *per se* that drives the FBR.

It is well established that the physical and chemical properties of materials modulate their ability to acquire a protein coat (26, 74) and this, in turn, can alter the extent of macrophage fusion and macrophage/FBGC phenotype (11, 75, 76–78). This concept has been mainly explored *in vitro* studies; however, implantation in mammals of materials with diverse surface properties elicits a very similar FBR and results in an essentially identical healing response (2, 16). It is possible that almost unavoidable adsorption of fibrinogen on the surface of implanted materials and its rapid conversion into fibrin polymer would largely negate the unique surface properties of materials. Hence, macrophages will likely interact with fibrin-coated material rather than directly with the material. The physical and adhesive properties of fibrin matrices deposited on the surface of various materials implanted in different locations and their ability to support FBGC formation remain to be elucidated. Nonetheless, given the role of fibrin polymer in promoting macrophage fusion, inhibition of thrombin may be a useful strategy to control the adverse effects of FBGCs. Furthermore, inhibition of fibrin formation leading to the reduced density of collagen in the fibrous capsule may be especially beneficial for certain medical devices, including biosensors and controlled drug delivery systems that often fail due to the formation of a diffusion barrier.

## MATERIALS AND METHODS

### Reagents

The hybridoma producing mouse mAb 59D8, which recognizes the N-terminal end of the β-chain of human and mouse fibrin was previously described (79). The mAb was purified using Protein A agarose and characterized by ELISA as previously described (80). The rat mAb M1/70 which recognizes mouse CD11b/CD18 (integrin Mac-1) was purified from conditioned media of hybridoma cells obtained from the American Tissue Culture Collection (Manassas, VA) and then conjugated to Alexa Fluor 488 (catalog number A20181) according to the manufacturer’s instructions (Thermo Fisher Scientific, Walther, MA). The rabbit polyclonal antibody (catalog number PA5-29734), which recognizes mouse fibrinogen was purchased from Thermo Fisher Scientific (Waltham, MA) and the rabbit polyclonal antibody directed against the mouse fibrinopeptide A (catalog number ab103648) was from Abcam (Cambridge, MA). The mouse anti-collagen 1α (catalog number sc-293182) and anti-elastin (catalog number sc-58756) mAbs were from Santa Cruz Biotechnology (Dallas, TX). The mouse anti-α -SMA mAb (catalog number MAB1420-SP) was from R&D systems. The rat anti-CD68 mAb (catalog number 14-0681-82) and the rabbit polyclonal anti-mouse collagen I (catalog number PA5-95137) were from Invitrogen (Carlsbad, CA). The mouse anti-fibrillin-1 mAb (catalog number MAB2502) was purchased from Millipore (Temecula, CA). The secondary antibodies, Alexa Fluor 647-conjugated goat anti-rabbit IgG and Alexa Fluor 488-conjugated goat anti-mouse IgG, were obtained from Invitrogen (Carlsbad, CA). Thrombin inhibitor argatroban monohydrate (catalog number A0487) was from Sigma Aldrich (St. Louis, MO). The protease inhibitor cocktail was purchased from Thermo Fisher Scientific (Walther, MA). Mouse fibrinogen (catalog number ab92791) was obtained from Abcam (Cambridge, MA) or purified from freshly isolated mouse blood. Fibrin-monomer was prepared by dissolving the fibrin clot in 20 mM acetic acid as previously described (81).

### Mice

Wild type (WT) C57BL/6J mice were purchased from The Jackson Laboratory (Bar Harbor, ME). Fibrinogen-deficient (Fg^-/-^) and fibrin-deficient (Fib^AEK^) mice were previously described (34, 82). All animals were given *ad libitum* access to food and water and maintained at 22 °C on a 12-h light/dark cycle.

### Thioglycollate-induced peritonitis

Eight-to twelve-week-old male and female mice were used in all experiments with age-and sex-matched WT and Fg^-/-^ animals selected for side-by-side comparison. Peritonitis in mice was induced by the intraperitoneal injection of 0.5 ml of a 4% Brewer thioglycollate (TG) solution (Sigma-Aldrich, St. Louis, MO) as described (83). Cells were collected 3 days after TG injection by peritoneal lavage with 5 ml ice-cold PBS with 5 mM EDTA. The total number of cells in the lavage fluid was counted using a hemacytometer.

### Biomaterial implantation and analyses of the retrieved explants

As an *in vivo* model for assessing the FBR to biomaterials, a well-characterized (27, 84) intraperitoneal implantation model was used. The peritoneal cavity provides a good site for studying the cell reactions caused by implantation because of the minimal contact with the interstitium of the normal tissue. Sections (1.5×0.5 cm) of sterile polychlorotrifluoroethylene (PCTFE) or polytetrafluoroethylene (PTFE) were implanted into the peritoneum of mice as previously described (50). Ten- to twelve-week old male and female mice were used in all experiments with age- and sex-matched wild-type and deficient animals selected for side-by-side comparison. Animals were humanely sacrificed 3, 7, and 14 days later, and retrieved explants were divided into two parts. One part was used for calculating the fusion index by immunofluorescence and the other part was used for histological analyses. The fibrinous material that covered the explant (capsule) retrieved from WT and Fib^AEK^ mice was carefully removed to expose the surface and saved for analyses of the protein composition by Western blotting. Before explantation, 2 ml of ice-cold PBS containing 5 mM EDTA was aseptically injected into the peritoneum, cells in the peritoneum were collected by lavage and counted. The percentage of macrophages in the lavage was determined by differential analysis of cytospin preparations dyed with Wright stain. The material deposited around the implant in the form of a fibrinous capsule was collected for Western blot, histological, ultrastructural, and AFM analyses.

### Preparation of plasma- and protein-coated PCTFE surfaces

The plasma for coating of PCTFE surfaces and isolation of fibrinogen was prepared from blood isolated from WT and Fg^-/-^ mice by cardiac puncture. Blood (0.3-0.6 ml) was drawn from each mouse using a 23G needle and insulin syringe. Anticoagulant citrate/dextrose solution was added to the blood at a 1:7 ratio. The blood was centrifuged at 3000 g for 15 min and the plasma was purified from the potential endotoxin contamination using high capacity endotoxin removal spin columns (ThermoFisher Scientific, Waltham, MA; 88274). The isolated plasma, purified mouse fibrinogen, and fibrin-monomer used in the “rescue” experiments were tested for endotoxin using Pierce™ Chromogenic Endotoxin Quant kit (ThermoFisher Scientific; A39552S) and the results showed that proteins contained < 0.7 EU/ml.

### Histological analyses

Retrieved implants with a surrounding capsule were fixed in 10% formalin for 24 h at 22 °C. After fixation, the materials were embedded in paraffin, sectioned, and stained according to a standard histological H&E procedure. The sections were mounted on a cover glass and imaged using the EVOS FL Auto (Thermo Scientific, Waltham, MA) wide-field microscope and a 40x objective.

### Isolation of cells from the capsule

On days 7 and 14 after surgery, the PCTFE sections were explanted from the mouse peritoneum. The capsule was removed and placed into a 35-mm dish (FluoroDish™; World Precision Instruments, Sarasota, FL) filled with the warm DMEM/F-12 medium for 24 hours in a cell culture incubator. During this time the majority of the cells migrated out of the cap. The remaining cells in the cap were removed by incubating the cap in a collagenase D solution (1 mg/ml) for 2 hours at 37 °C followed by collecting the cells by centrifugation at 300 x g for 3 minutes. The cells were resuspended in DMEM/F-12 and added to the initial pool of cells in a Fluorodish. Adherent cells were fixed with 2% formaldehyde in PBS for 30 minutes, permeabilized with 0.1% Triton X-100 for 30 minutes, blocked with 1% BSA for 1 hour at 22 °C, and used for immunofluorescence. Macrophages were separated from fibroblasts with the use of the EasySep Mouse selection kit (StemCell Technologies, Vancouver, BC, Canada) with mAb F4/80 conjugated to phycoerythrin.

### Immunofluorescence

To determine the fusion index, retrieved PCTFE sections were fixed with 2% formaldehyde in PBS for 30 min at 22 °C. Adherent cells were permeabilized with 0.1% Triton X-100 in PBS for 30 min and then washed three times with PBS containing 1% BSA. Nuclei were labeled with DAPI and actin was labeled with Alexa Fluor 488-conjugated phalloidin according to the manufacturer’s recommendation (Thermo Scientific, Waltham, MA). Samples were mounted in Prolong Diamond (Thermo Scientific, Waltham, MA) and imaged with a Leica SP5 or Leica SP8 laser scanning confocal microscopes using 40x/1.3 NA oil objective. The fusion index was determined as previously described (85) and is defined as the fraction of nuclei within FBGCs expressed as the percentage of the total nuclei counted. Five to six fields imaged by a 40×objective that contained ∼100–200 cells were analyzed for each experimental condition. The macrophage spreading was assessed using NIH ImageJ software by quantitating the surface area (µm^2^) of adherent macrophages.

To examine the presence of various extracellular matrix proteins in the capsule, paraffin blocks were cut and the sections deparaffinized followed by dehydration in ethanol. After incubation in PBS+1% BSA, the sections were incubated with primary antibodies (anti-fibrinogen, anti-fibrin, anti-α-SMA, anti-CD68, anti-collagen 1α, anti-fibrillin, and anti-elastin) for 1 h at 22 °C followed by secondary antibodies conjugated to Alexa Fluor 647. In addition to the abovementioned primary antibodies, cells isolated from the capsule were incubated with mAb M1/70 directed to a mouse integrin Mac-1. Samples were mounted in Prolong Diamond (Thermo Scientific) and imaged with a 40x oil objective using an SP5 confocal microscope.

### Western blotting

The implants inserted into WT, Fg^-/-^ and Fib^AEK^ mice were explanted after 3, 7 and 14 days and the fibrinous capsule was removed from the surface (it is easily detachable with no adhesive resistance). The retrieved material sections and fibrinous capsules were placed in PBS containing protease inhibitors and then the loading buffer containing SDS (1% final concentration), 8 M urea, and 10 mM N-ethylmaleimide was added. Samples were electrophoresed on 7.5% SDS-polyacrylamide gels and proteins were transferred onto the Immobilon-P membrane (Millipore, New Bedford, MA). Blots were probed with anti-fibrinogen polyclonal antibody (1:100,000 dilution), anti-fibrinopeptide A polyclonal antibody (1:1000 dilution) and fibrin-specific mAb 59D8 (1 µg/ml) followed by goat anti-rabbit and rabbit anti-mouse secondary antibodies conjugated with horseradish peroxidase (1:10,000 dilution). Bound antibodies were detected by reaction with a SuperSignal West Pico Chemiluminescent Substrate (Thermo Scientific, Grand Island, NY; Cat. No 34577).

### TEM analysis of the capsule

1.5 cm x 0.5 cm PCTFE pieces implanted into mouse peritoneum were explanted after 14 days, fixed with 2.5% glutaraldehyde in 0.1 M PBS (pH 7.4) at 4 ^°^C overnight, and then treated with 1% OsO_4_ in 0.1 M PBS for 1 h. Subsequently, cells were washed with 0.1 M PBS and then dehydrated using acetone. Finally, the implants with the surrounding capsule were flat-embedded into Spur’s EPOXY Resin, and 70-nm sections were obtained by longitudinal or vertical sectioning of the sample. Sections were post-stained with uranyl acetate and Sato’s lead citrate. Micrographs were taken using a Philips CM 12 TEM with a Gatan Model 791 camera.

### Atomic Force Microscopy (AFM)

An Asylum Research MFP-3D-BIO AFM was used to conduct the force-indentation measurements. AFM probes with sphere-cone geometry were used (LRCH-750 Team NanoTec, Villingen-Schwenningen, Germany). The spring constants (nominal k∼0.2 N.m^-1^) were determined using the thermal energy dissipation method built-in the Asylum Research image acquisition software. Samples were measured at room temperature in DPBS. Quasi-static measurements with a cantilever approach and retraction speed 2 µm s^-1^ were conducted to collect elastic modulus data. In areas in the center of the samples, 15 grids of 5×4 indentations were acquired by applying a trigger force of 45 nN which resulted in 10-15µm of indentation. The force-indentation curves were fitted to a quasi-static contact model for a sphero-conical indenter (86, 87).

### RT-PCR analysis of collagen I expression in macrophages isolated from the capsule

PCTFE pieces were implanted into the mouse peritoneum for 14 days. The total population of cells was isolated from the capsule and macrophages were separated from fibroblasts using by magnetic beads. Total RNA was extracted using RNeasy Plus mini kit (Qiagen, Hilden, Germany; Cat. No 74134) and resuspended in 20 ml of RNase-free water supplemented with 0.1 mM EDTA. 1 µg of total RNA was used to generate cDNA using SuperScript III Reverse Transcriptase (Invitrogen). PCR was performed with the generated cDNA and premixed 2× Taq polymerase solution (Promega) in an MJ Mini Thermal Cycler (BioRad). The primers for PCR analyses were purchased from Integrated DNA Technologies (Iowa, USA) and included GTCCCAGTGGTCCTCCCGGTC-3’ (*forward*) and 5’-CAGGGGGACCAGCCAATCCAG-3’ (*reverse*) primers. WI-38 fibroblasts cultured in DMEM F12 media supplemented with glutamine were used as a control cell line expressing collagen type I.

### Statistical analyses

Unless indicated otherwise results are shown as the mean ± S.D. from three independent experiments. Samples that passed the normal distribution test were analyzed by a two-tailed t-test. The remaining samples were analyzed by the Mann-Whitney U test. Data were considered significantly different if p < 0.05. The statistical analysis was performed using GraphPad Instat software.

### Study approval

All animal studies comply with the National Institutes of Health guide for the care and use of laboratory animals. Experiments were performed according to animal protocols approved by the Institutional Animal Care and Use Committees at Arizona State University.

## Supporting information

Supplemental material all

## Author Contributions

A.B., N.P.P., J.K., D.L., A.Z. performed experiments and analyzed data. M.J.F. and P.B. provided tools and reagents. T.P.U. and M.J.F. conceived the project. T.P.U., A.B., M.J.F., and R.R. designed experiments, analyzed data, and wrote the manuscript.

## Acknowledgment

This work was supported by the National Institutes of Health grants HL63199 (TU) and DK112778 (MJF). We acknowledge the use of instruments within the W.M. Keck Bioimaging Facility at Arizona State University. Image data were collected using a Leica TCS SP5 LSCM (the National Institutes of Health SIG award S10 RR027154) and Leica TCS SP8 LSCM (the NIH SIG award S10 OD023691).

## Conflict of Interest Statement

The authors have declared that no conflict of interest exists.

## SUPPLEMENTAL FIGURES

**Supplemental figure 1. The number of cells in the inflamed peritoneum of WT and Fg^-/-^ mice.** Thioglycollate solution was injected into the mouse peritoneum and peritoneum lavage was collected after 72 hours. Results shown are mean ± SD from three independent experiments (n=3 WT and n=3 Fg^-/-^ mice). ns, not significant.

**Supplemental figure 2. The FBR to implanted PTFE biomaterials.** (A) PTFE sections were implanted in the peritoneum of wild type and Fg^-/-^ mice for 7 days and explants were separated from the surrounding fibrinous capsule. Since PTFE plastic is nontransparent and consequently unamenable to immunocytochemistry, firmly adherent cells were removed from the surface using a cell scraper, centrifuged, and resuspended in DMEM/F-12, Cells were allowed to adhere to polylysine-coated coverslips and incubated with Alexa Fluor 546-conjugated phalloidin (white) and DAPI (teal). Representative confocal images are shown. FBGCs are outlined (yellow). The scale bar is 50 µm. (B) Macrophage fusion was assessed as a fusion index. Five to six random 20× fields were used per sample to count nuclei. (C) Explants were fixed, paraffin-embedded, sectioned, and stained according to a standard H&E method. Representative images of stained cross-sections are shown. The scale bar is 50 µm. (D) The thickness of granulation tissue around the implants retrieved from WT and Fg^-/-^ mice was determined using ImageJ software. Ten random fields were used per sample to measure the thickness of cross-sections. Results shown are mean ± SD from two independent experiments (n=2 WT and n=2 Fg^-/-^ mice). Mann-Whitney *U* test was used to calculate significance \*\*\**p* < .001

**Supplemental figure 3. Analyses of multinucleation and the granulation tissue formation of implants precoated with plasma.** PCFTE sections pre-coated with mouse plasma were implanted into Fg^-/-^ mice for 3 days. Uncoated sections implanted in WT and Fg^-/-^ mice served as controls. (A) Explants were separated from the surrounding fibrous capsule, fixed, and incubated with Alexa Fluor 546-conjugated phalloidin (white) and DAPI (teal). FBGCs are outlined (yellow). Representative confocal images are shown. (B) Representative images of the granulation tissue formed around different implants. (C) Fusion indices were determined as described in Materials and Methods. (D) The number of nuclei per FBGC was analyzed to assess the extent of multinucleation. 30 FBGCs were analyzed from 6-8 fields. (E) The thickness of the granulation tissue formed around uncoated and precoated implants shown in B. The scale bar is 50 µm. Results shown are mean ± S.D. of four independent experiments (n=4 WT and n=4 Fg^-/-^ mice). Two-tailed *t-*test and Mann-Whitney *U* test were used to calculate significance. ns, not significant, *p <* .05, \*\**p* < .01, \*\*\**p* < .001

**Supplemental figure 4.** Effect of argatroban on the total number of cells adherent to the surface of PCTFE biomaterials implanted in WT mice. Argatroban (9 mg/kg and 18 mg/kg) was injected i.p. for 5 days before implantation of materials and subsequently for 7 days post-surgery. Control mice were injected with PBS. The number of cells on the surface of PCTFE explants retrieved from control and argatroban-treated mice was determined by counting nuclei. Results shown are mean ± SD from three independent experiments (6 mice per group). Mann-Whitney *U* test was used to calculate significance. ns, not significant, \*\**p* < .01

**Supplemental figure 5. The number of cells in lavage obtained from WT and Fib^AEK^ mice.** The peritoneal cells were collected from WT and Fib^AEK^ mice before the explantation of the PCTFE implants at various time points. Results shown are mean ± S.D. of four independent experiments. n=5-6 (WT) and n=5 (Fib^AEK^) mice. Mann-Whitney *U* test was used to calculate significance. ns, not significant, \*\**p* < .01.

**Supplemental figure 6. Western blot analysis of fibrin(ogen) species deposited on the surface and in the capsule formed around PCTFE sections implanted into Fib^AEK^ mice.** (A) The specificity of antibodies recognizing total fibrinogen (anti-total Fg; 1: 50,000 dilution) and fibrinopeptide A (anti-FpA; 1:2000 dilution). (A) Purified mouse fibrinogen from WT and Fib^AEK^ mice were electrophoresed on 7.5% polyacrylamide gel followed by Western blotting using fibrinogen- and FpA-specific antibodies. (B) Analysis of surfaces retrieved 3, 7, and 14 days after implantation. The fibrinous capsule was removed from the explanted material and exposed sections were placed into PBS containing protease inhibitors followed by the SDS-PAGE loading buffer and Western blotting using anti-total Fg, anti-FpA, and anti-fibrin antibodies. The arrowhead in the right panel shows the IgG-reactive product. The data shown are representative of samples obtained from four mice. (C) Analysis of fibrinogen species present in the capsule formed at different time points.

**Supplemental figure 7. Deposition of fibrillin and elastin in the capsule formed around the implants in WT and Fib^AEK^ mice.** PCTFE sections were implanted in WT and Fib^AEK^ mice for 3, 7, and 14 days and the capsules formed were analyzed for the presence of fibrillin and elastin. (A and C) Representative immunofluorescence images of the samples incubated with antibodies that recognize fibrillin. (B and D) Representative immunofluorescence images of the samples incubated with antibodies that recognize elastin are shown. The scale bars are 50 µm. (E and F) Quantification of fluorescence intensities of images of fibrillin and elastin deposition. Results shown are mean ± S.D. from the areas of 0.2 mm^2^; n=100 for each sample. A two-tailed t-test was used to calculate significance. ns, not significant, \*\**p* < .01

**Supplemental figure 8.** Detection of collagen expression in macrophages isolated from the fibrous capsule. (A) PCTFE sections were implanted in WT mice for 7 days and the cells in the capsule were isolated as described in Materials and Methods. The cells were allowed to adhere to the surface of a FluoroDish, fixed and incubated with anti-CD11b mAb M1/70 and anti-collagen polyclonal antibodies followed by Alexa Fluor 488- and Alexa Fluor 647-conjugated secondary antibodies. Cells also were stained with Alexa Fluor 568-conjugated phalloidin and DAPI. The scale bar is 15 µm. (B) Detection of mRNA for collagen I in macrophages isolated from the capsule using RT-PCR. PCTFE sections were implanted in WT mice for 14 days and the cells in the capsule were isolated as described in Materials and Methods. Macrophages were separated from fibroblasts using mAb F4/80-coated magnetic beads. The PCR analysis was performed as described in Materials and Methods. WI-38 fibroblasts served as a control.

**Supplemental figure 9. Macrophages express fibrillin (A) and elastin (B)**. Cells isolated from a 14-day capsule formed in WT mice were allowed to adhere to the surface of a FluoroDish, fixed, and incubated with primary antibodies (anti-CD11b mAb M1/70, anti-fibrillin, and anti-elastin) followed by a secondary antibody.

## Notes

### Competing Interest Statement

The authors have declared no competing interest.

