## Supplemental material all for "Fibrin Polymer on the Surface of Biomaterial Implants Drives the Foreign Body Reaction"

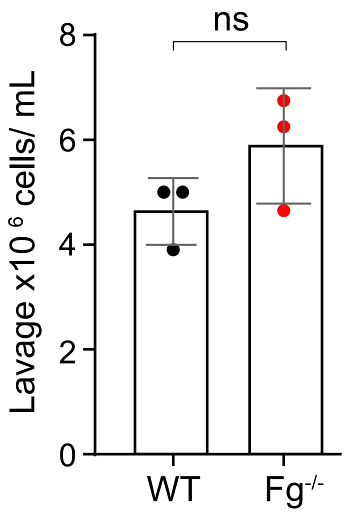


**Supplemental figure 1. The number of cells in the inflamed peritoneum of WT and Fg$^{-/-}$ mice.** Thioglycollate solution was injected into the mouse peritoneum and peritoneum lavage was collected after 72 hours. Results shown are mean ± SD from three independent experiments (n=3 WT and n=3 Fg$^{-/-}$ mice). ns, not significant

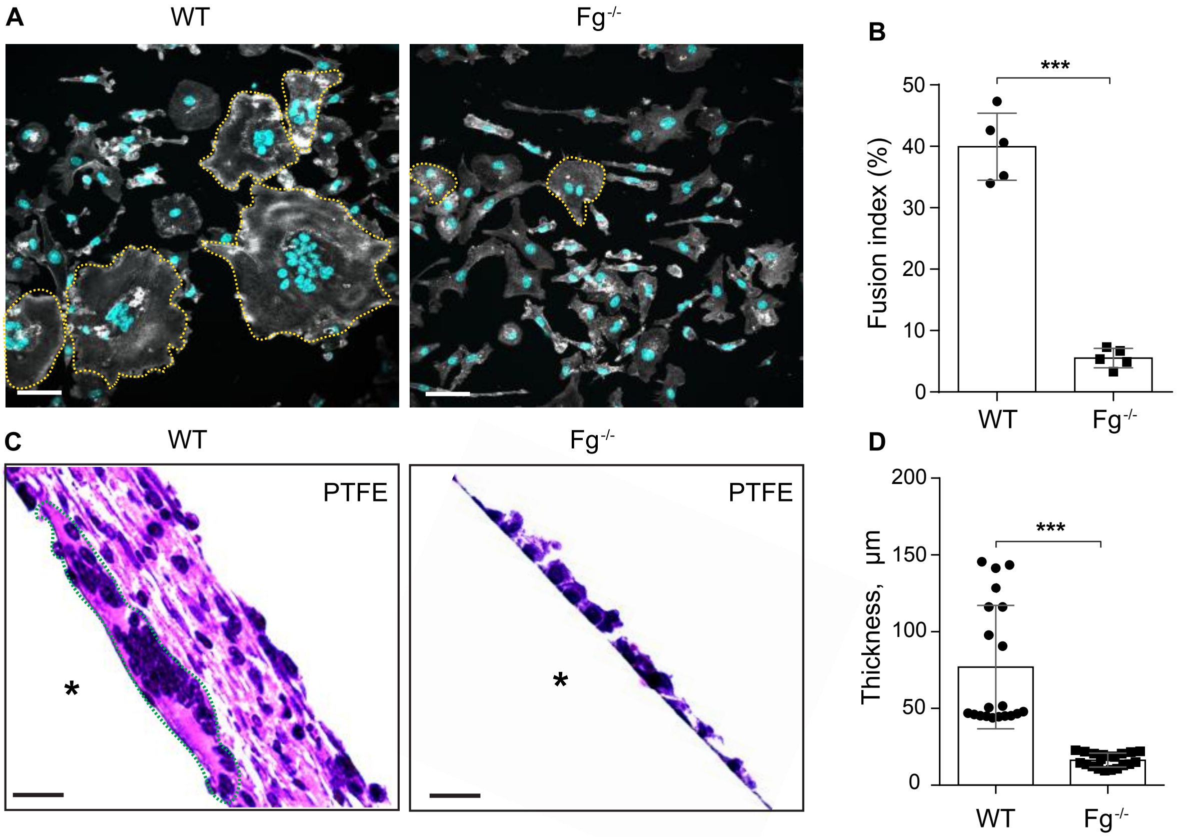


**Supplemental figure 2. The FBR to implanted PTFE biomaterials. (A)** PTFE sections were implanted in the peritoneum of wild type and $Fg^{-/-}$ mice for 7 days and explants were separated from the surrounding fibrinous capsule. Since PTFE plastic is nontransparent and consequently unamenable to immunocytochemistry, firmly adherent cells were removed from the surface using a cell scraper, centrifuged, and resuspended in DMEM/F-12, Cells were allowed to adhere to polylysine-coated coverslips and incubated with Alexa Fluor 546-conjugated phalloidin (white) and DAPI (teal). Representative confocal images are shown. FBGCs are outlined (yellow). The scale bar is 50 µm. **(B)** Macrophage fusion was assessed as a fusion index. Five to six random 20× fields were used per sample to count nuclei. **(C)** Explants were fixed, paraffin-embedded, sectioned, and stained according to a standard H&E method. Representative images of stained cross-sections are shown. The scale bar is 50 µm. **(D)** The thickness of granulation tissue around the implants retrieved from WT and $Fg^{-/-}$ mice was determined using ImageJ software. Ten random fields were used per sample to measure the thickness of cross-sections. Results shown are mean ± SD from two independent experiments (n=2 WT and n=2 $Fg^{-/-}$ mice). Mann-Whitney *U* test was used to calculate significance ***$p < .001$

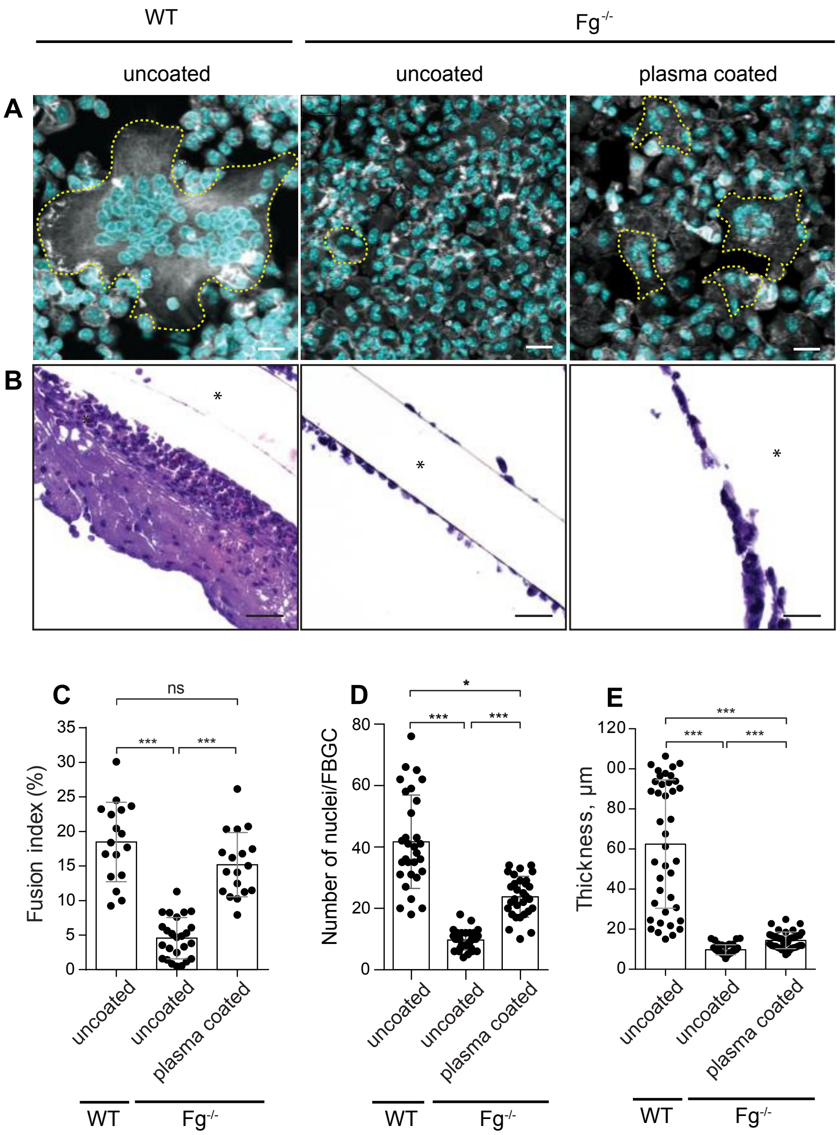


**Supplemental figure 3. Analyses of multinucleation and the granulation tissue formation of implants precoated with plasma.** PCFTE sections pre-coated with mouse plasma were implanted into Fg$^{-/-}$ mice for 3 days. Uncoated sections implanted in WT and Fg$^{-/-}$ mice served as controls. **(A)** Explants were separated from the surrounding fibrous capsule, fixed, and incubated with Alexa Fluor 546-conjugated phalloidin (white) and DAPI (teal). FBGCs are outlined (yellow). Representative confocal images are shown. **(B)** Representative images of the granulation tissue formed around different implants. **(C)** Fusion indices were determined as described in Materials and Methods. **(D)** The number of nuclei per FBGC was analyzed to assess the extent of multinucleation. 30 FBGCs were analyzed from 6-8 fields. **(E)** The thickness of the granulation tissue formed around uncoated and precoated implants shown in B. The scale bar is 50 μm. Results shown are mean ± S.D. of four independent experiments (n=4 WT and n=4 Fg$^{-/-}$ mice). Two-tailed t-test and Mann-Whitney *U* test were used to calculate significance. ns, not significant, p < .05, **p < .01, ***p < .001

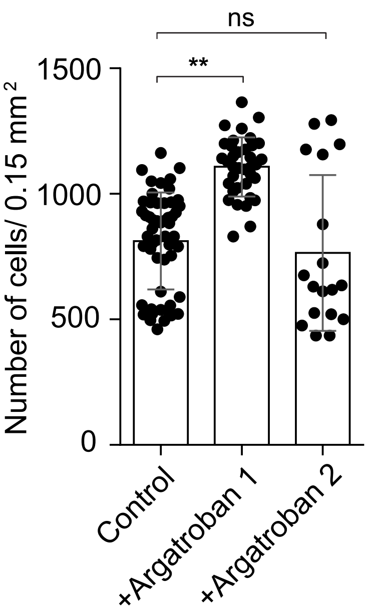


**Supplemental figure 4. Effect of argatroban on the total number of cells adherent to the surface of PCTFE biomaterials implanted in WT mice.** Argatroban (9 mg/kg and 18 mg/kg) was injected i.p. for 5 days before implantation of materials and subsequently for 7 days post-surgery. Control mice were injected with PBS. The number of cells on the surface of PCTFE explants retrieved from control and argatroban-treated mice was determined by counting nuclei. Results shown are mean ± SD from three independent experiments (6 mice per group). Mann-Whitney *U* test was used to calculate significance. ns, not significant, **$p < .01$

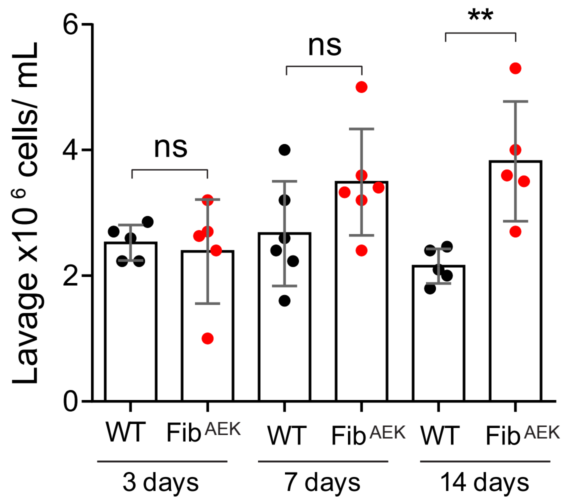


**Supplemental figure 5. The number of cells in lavage obtained from WT and Fib$^{AEK}$ mice.** The peritoneal cells were collected from WT and Fib$^{AEK}$ mice before the explantation of the PCTFE implants at various time points. Results shown are mean ± S.D. of four independent experiments. n=5-6 (WT) and n=5 (Fib$^{AEK}$) mice. Mann-Whitney *U* test was used to calculate significance. ns, not significant, **p < .01

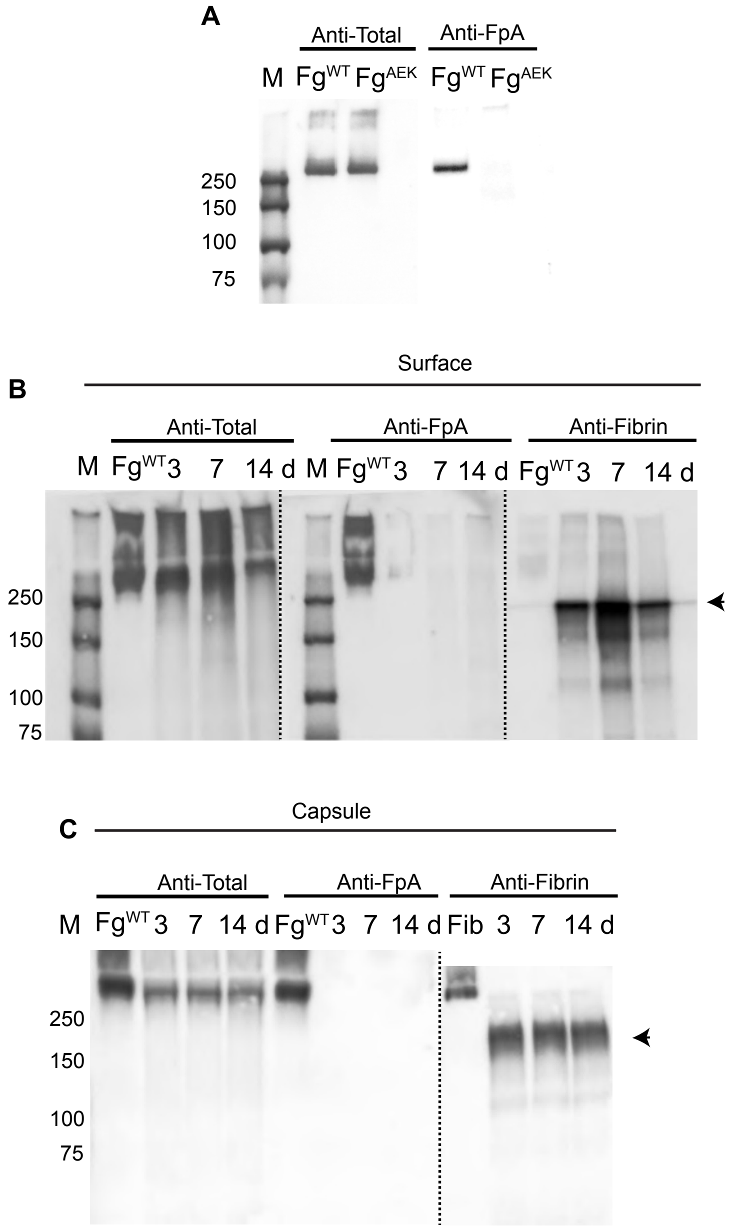


**Supplemental figure 6. Western blot analysis of fibrin(ogen) species deposited on the surface and in the capsule formed around PCTFE sections implanted into Fib$^{AEK}$ mice.** The specificity of antibodies recognizing total fibrinogen (anti-total Fg; 1: 50,000 dilution) and fibrinopeptide A (anti-FpA; 1:2000 dilution). **(A)** Purified mouse fibrinogen from WT and Fib$^{AEK}$ mice were electrophoresed on 7.5% polyacrylamide gel followed by Western blotting using fibrinogen- and FpA-specific antibodies. **(B)** Analysis of surfaces retrieved 3, 7, and 14 days after implantation. The fibrinous capsule was removed from the explanted material and exposed sections were placed into PBS containing protease inhibitors followed by the SDS-PAGE loading buffer and Western blotting using anti-total Fg, anti-FpA, and anti-fibrin antibodies. The arrowhead in the right panel shows the IgG-reactive product. The data shown are representative of samples obtained from four mice. **(C)** Analysis of fibrinogen species present in the capsule formed at different time points

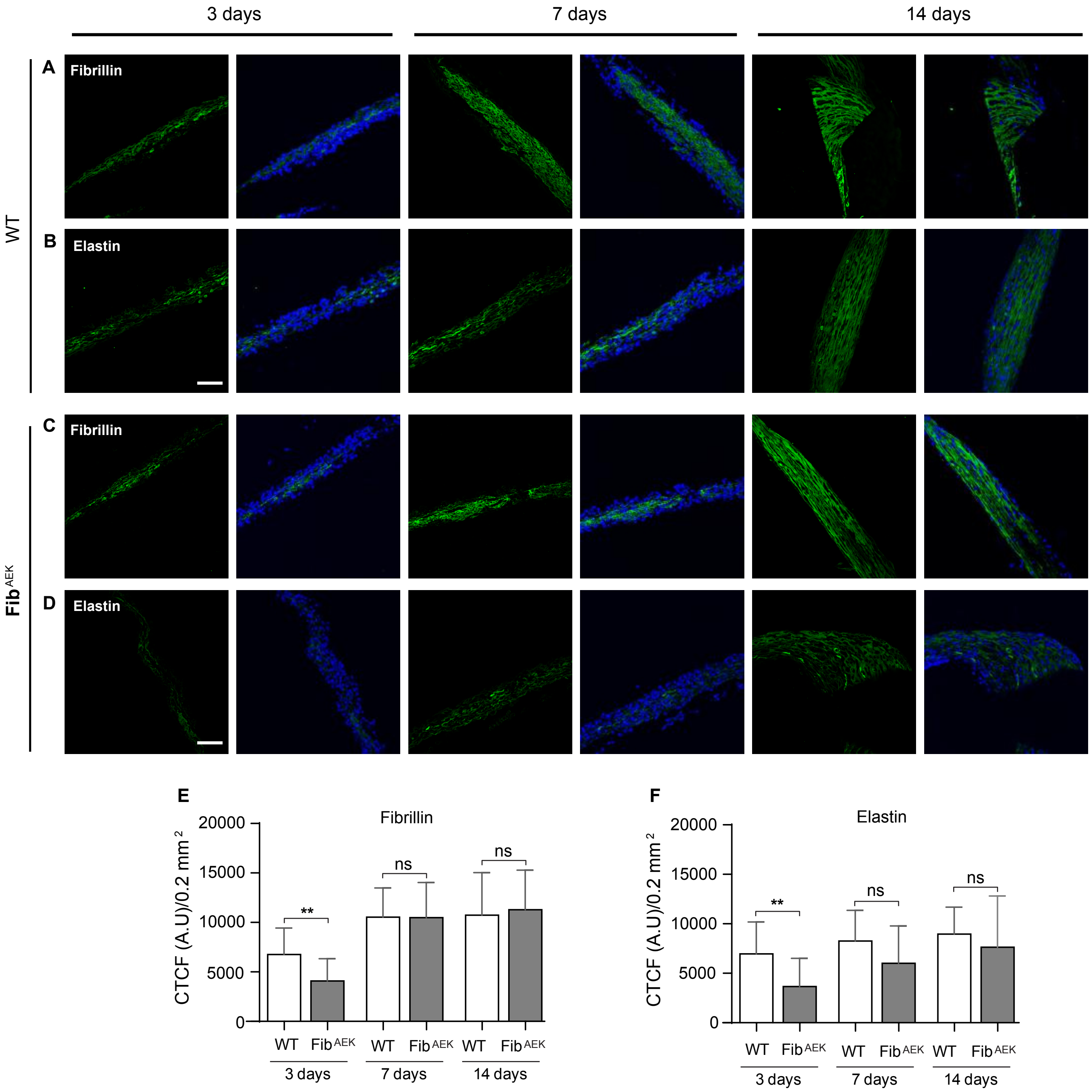


**Supplemental figure 7. Deposition of fibrillin and elastin in the capsule formed around the implants in WT and Fib$^{AEK}$ mice.** PCTFE sections were implanted in WT and Fib$^{AEK}$ mice for 3, 7, and 14 days and the capsules formed were analyzed for the presence of fibrillin and elastin. **(A** and **C)** Representative immunofluorescence images of the samples incubated with antibodies that recognize fibrillin. **(B** and **D)** Representative immunofluorescence images of the samples incubated with antibodies that recognize elastin are shown. The scale bars are 50 µm. **(E** and **F)** Quantification of fluorescence intensities of images of fibrillin and elastin deposition. Results shown are mean ± S.D. from the areas of 0.2 mm$^2$; n=100 for each sample. Two-tailed t-test was used to calculate significance. ns, not significant, **p < .01

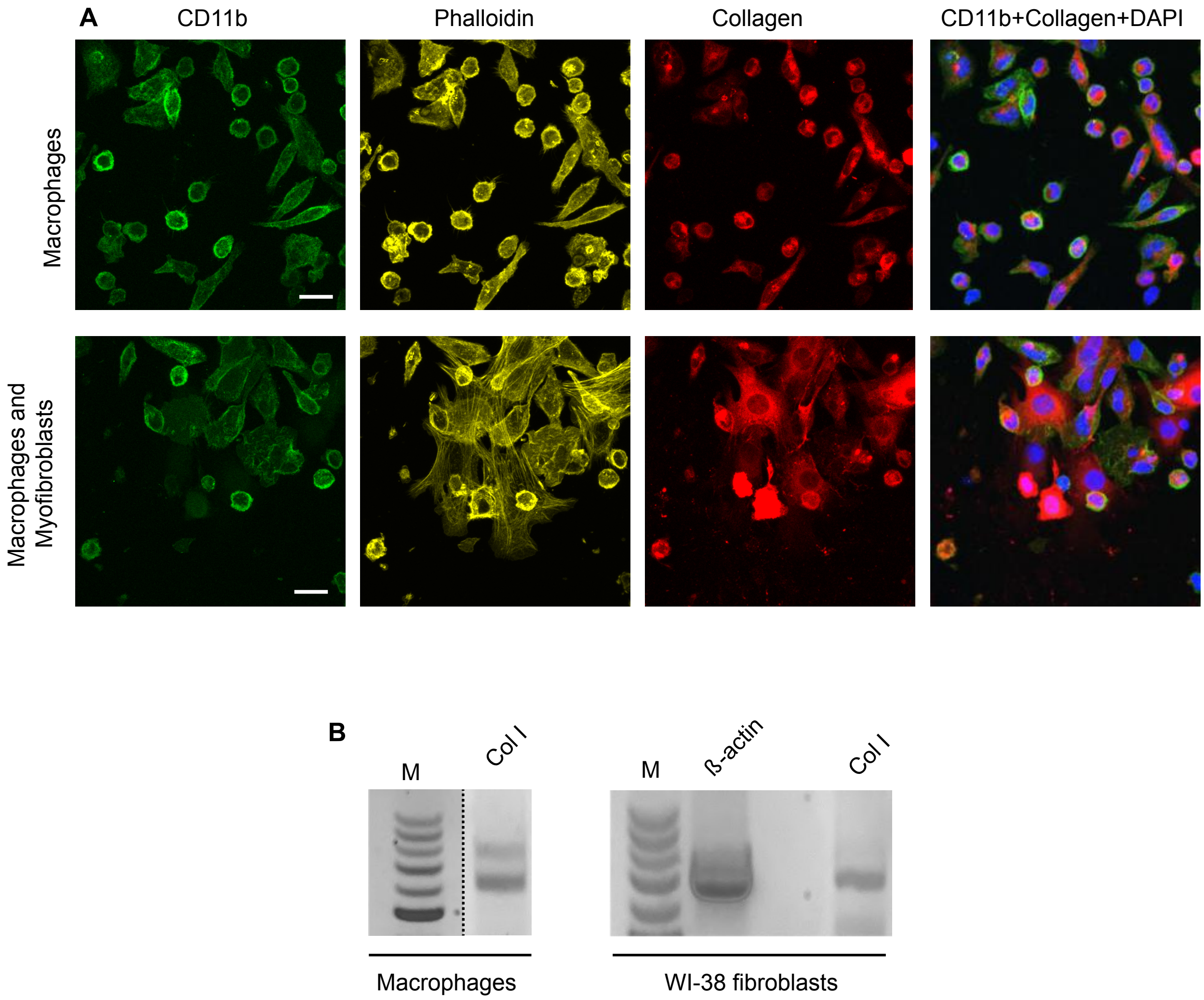


**Supplemental figure 8. Detection of collagen expression in macrophages isolated from the fibrous capsule. (A)** PCTFE sections were implanted in WT mice for 7 days and the cells in the capsule were isolated as described in Materials and Methods. The cells were allowed to adhere to the surface of a FluoroDish, fixed and incubated with anti-CD11b mAb M1/70 and anti-collagen polyclonal antibodies followed by Alexa Fluor 488- and Alexa Fluor 647-conjugated secondary antibodies. Cells also were stained with Alexa Fluor 568-conjugated phalloidin and DAPI. The scale bar is 15 µm. **(B)** Detection of mRNA for collagen I in macrophages isolated from the capsule using RT-PCR. PCTFE sections were implanted in WT mice for 14 days and the cells in the capsule were isolated as described in Materials and Methods. Macrophages were separated from fibroblasts using magnetic beads. The PCR analysis was performed as described in Materials and Methods. WI-38 fibroblasts served as a control

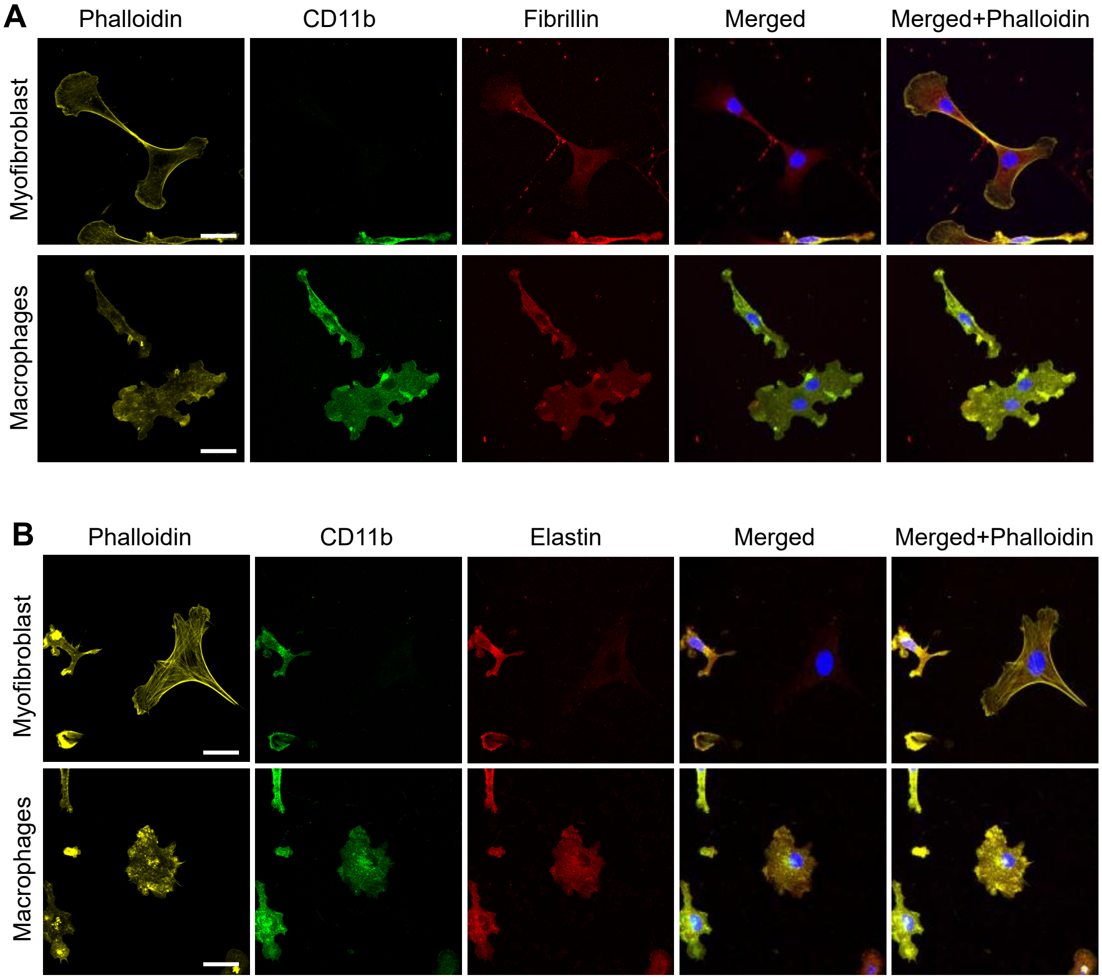


**Supplemental figure 9. Macrophages express fibrillin (A) and elastin (B).** Cells isolated from a 14-day capsule formed in WT mice were allowed to adhere to the surface of a FluoroDish, fixed, and incubated with primary antibodies (anti-CD11b mAb M1/70, anti-fibrillin, and anti-elastin) followed by a secondary antibody. The scale bar is 15 μm.
